# A Multiscale Signaling–Biophysical Framework Reveals Mechanisms of Macrophage-Mediated RBC Clearance in Sickle Cell and Gaucher Disease

**DOI:** 10.64898/2026.04.20.719505

**Authors:** Zhaojie Chai, Nazanin Ahmadi Daryakenari, George Em Karniadakis

**Author notes:** To whom correspondence should be addressed: george. Zhaojie Chai and Nazanin Ahmadi Daryakenari contributed equally to this work.

## Abstract

Red blood cell (RBC) clearance by macrophages maintains blood homeostasis and is dysregulated in the hemolytic disorder sickle cell disease (SCD) and the lysosomal storage disorder Gaucher disease (GD), where biophysical and biochemical alterations promote premature phagocytosis. We develop a multiscale hybrid modeling framework integrating signaling dynamics, biophysical simulations, and machine learning to investigate the mechanisms governing RBC phagocytosis in these diseases. Our approach couples a systems biology model of macrophage–RBC signaling with Dissipative Particle Dynamics (DPD) simulations of molecular diffusion and membrane interactions, and leverages Physics-Informed Neural Networks (PINNs) for robust parameter inference. The DPD framework provides mechanistic insight into antibody diffusion, receptor engagement, and membrane-level interactions during macrophage–RBC contact, generating spatially resolved trajectories of CD47–SIRP*α* signaling and antibody–receptor binding that serve as intermediate observables constraining the signaling model. The model accurately captures differential phagocytic responses between healthy and altered RBCs, revealing diminished inhibitory signaling and changes in SHP1-mediated pathways in both SCD and GD. Identifiability analysis combining Fisher Information Matrix diagnostics and profile likelihood confirms that parameters governing the CD47–SIRP*α*–SHP1 axis are among the most robustly recoverable, and simulations of therapeutic perturbations with anti-SIRP*α* antibodies demonstrate modulation of engulfment outcomes. We further employ Physics-Informed Kolmogorov-Arnold Networks (PIKANs) as an alternative to standard PINNs, demonstrating improved robustness under noise and sampling variability. More broadly, our multiscale platform linking biophysical simulation with systems-level inference is generalizable, offering mechanistic insights and computational tools for therapeutic exploration in diseases involving dysregulated phagocytosis.

**Significance statement:** Red blood cells are normally removed from circulation by macrophages through tightly regulated molecular signals. In diseases such as sickle cell disease and Gaucher disease, this clearance process becomes abnormal, contributing to anemia and other complications. However, the mechanisms linking the physical properties of red blood cells to immune signaling remain poorly understood. Here we develop a multiscale computational framework that combines particle-based biophysical simulations, systems biology models, and physics-informed machine learning. This approach provides a quantitative framework to interpret how changes in red blood cell mechanics and surface signaling disrupt the CD47-SIRP*α* inhibitory pathway that normally prevents phagocytosis. The framework provides a predictive platform for studying immune clearance and may help guide therapeutic strategies targeting red blood cell–macrophage interactions.

## Introduction

Red blood cells (RBCs) circulate for about 120 days, after which aged or damaged RBCs are removed by macrophages in the spleen to maintain healthy blood cell populations. This clearance process relies on a balance of “self” signals that prevent unwarranted phagocytosis and “eat-me” signals that mark defective RBCs for removal. A well-known inhibitory signal is the interaction between RBC surface protein CD47 and macrophage SIRP*α*, which engages a “don’t eat me” pathway (32; 80; 65; 73). Conversely, exposure of phosphatidylserine (PS) on RBCs or opsonization of RBCs can trigger pro-phagocytic pathways via macrophage Fc*γ* receptors and other receptors (7; 55). These molecular interactions converge on the macrophage cytoskeletal machinery. For instance, SIRP*α* signaling recruits the phosphatase SHP1, which in turn modulates myosin IIa activity required for engulfment (26). Through such mechanisms, macrophages can discriminate healthy from aberrant RBCs and execute phagocytosis appropriately.

In sickle cell disease and Gaucher disease, pathological alterations in RBC properties upset this balance. SCD, caused by a hemoglobin mutation, produces misshapen, rigid RBCs with altered membrane protein display. These sickle red blood cells tend to become trapped in the spleen and are cleared more rapidly than normal cells, which contributes to anemia and acute splenic sequestration crises (50). Studies by Khandelwal et al. (39) indicated that sickle cells may have reduced effective CD47-SIRP*α* signaling, diminishing their ability to convey a “self” signal and thereby permitting higher phagocytic uptake. In GD, deficiency of glucocerebrosidase leads to systemic accumulation of glucosylceramide, including altered lipid composition of RBC membranes, as shown by Franco et al. (28) and Dupuis et al. (21). Gaucher red blood cells exhibit abnormal biomechanics, including increased rigidity and greater aggregation propensity, together with altered membrane composition, which affects their deformability and surface signaling, as described by Adar and colleagues (2; 1). These alterations are thought to promote splenic trapping and phagocytosis of Gaucher cells, contributing to anemia and splenomegaly in patients (11).

Despite extensive characterization of red blood cell deformability and adhesion in both SCD and GD, the precise multiscale mechanisms by which these alterations affect macrophage signaling and engulfment, that is, erythrophagocytosis, remain incompletely understood. To address this knowledge gap, we developed an integrative computational model that merges biochemical signaling pathways with biophysical cell interaction dynamics. Because experimental *in vitro* measurements alone cannot fully resolve the spatial and temporal dynamics of macrophage–RBC interactions, we employ complementary computational approaches. Advances in computational power over the past two decades have stimulated the development and application of multiscale biophysical models for studying complex cellular systems (70; 10; 13). In parallel, machine learning approaches have increasingly been applied to collective cell dynamics and particle-based simulations (88; 87).

In our framework, a system of ordinary differential equations (ODEs) captures the key signaling network involved in RBC phagocytosis, including the dynamics of the CD47-SIRP*α*-SHP1 axis and downstream effectors such as myosin IIa. This ODE-based signaling model is informed by experimental data for specific molecular species, such as the kinetics of CD47-SIRP*α* binding (54). However, purely ODE-based formulations cannot directly capture spatial diffusion of molecules or the physical deformation and contact between a macrophage and a red blood cell. To address these limitations, we integrated a dissipative particle dynamics framework to model the macrophage microenvironment and the mesoscopic interaction between the RBC and macrophage. DPD is a particle-based simulation technique particularly well-suited for soft biological matter (25; 11). We model representative disease-induced perturbations rather than patient-specific parameterization. In our context, membranes of the macrophage and RBC are represented as collections of interacting particles, while signaling molecules are explicitly simulated as diffusing particles in the surrounding fluid. This formulation allows us to incorporate spatial and stochastic aspects of molecular transport and cell contact that influence downstream signaling. For example, snapshots from the DPD simulations illustrate an RBC and macrophage suspended in a fluid medium with diffusing signaling particles, highlighting how local concentrations evolve dynamically as the cells engage. We validated the DPD component by confirming that simulated molecules undergo random-walk trajectories and that the measured mean squared displacement increases linearly with time, yielding diffusion coefficients consistent with experimentally reported values for comparable biomolecules (58; 27). This validation supports the physical fidelity of the integrated environment and provides confidence that the hybrid ODE-DPD framework can generate biologically plausible constraints for downstream signaling inference. While the multiscale ODE–DPD framework provides mechanistic descriptions of membrane interactions and signaling dynamics, translating these simulations into quantitative predictions requires reliable estimation of kinetic parameters. In such models, parameter estimation is often hindered by both structural and practical non-identifiability. Structural non-identifiability refers to situations in which parameters cannot be uniquely recovered even under idealized, noise-free conditions given the model equations (16; 51). Practical non-identifiability, in contrast, arises from finite and noisy data, limited observables, or unsuitable experimental design, leading to parameter correlations or flat likelihood regions despite theoretical recoverability (33). In nonlinear dynamical systems such as the signaling model considered here, likelihood surfaces may exhibit extended ridges, parameter compensation, or strongly correlated directions (79; 31). Local sensitivity-based measures such as the Fisher Information Matrix (FIM) approximate uncertainty through curvature at the optimum but may provide misleading conclusions in the presence of nonlinear effects or poorly conditioned parameter directions (79). Similarly, bootstrap or Monte Carlo variance estimates may fail to reveal near-non-identifiable manifolds when multiple parameter combinations explain the data equally well (34).

To obtain a robust assessment, we combine sensitivity-based Fisher Information Matrix (FIM) analysis with profile likelihood, which systematically explores the likelihood surface through repeated re-optimization and enables the identification of finite confidence intervals and functional parameter dependencies (64; 79). These identifiability analyses directly inform parameter selection, search constraints, and the design of the subsequent physics-informed inverse modeling strategy.

Building on this identifiability-aware framework, we employ physics-informed neural networks to infer unknown kinetic parameters while enforcing consistency with the governing ODE system. PINNs enable simultaneous inference of system states and parameters by embedding the differential equations directly into the training objective through automatic differentiation, which is particularly advantageous under partial observation and noisy measurement regimes. Physics-informed learning has emerged as a powerful paradigm for solving forward and inverse problems in nonlinear dynamical systems by embedding governing equations directly into the loss function (62; 38). In biomedical and systems biology contexts, physics-informed learning has been successfully applied to gray-box identification (5; 61), inverse problems in systems biology and pharmacology (19; 4; 59), and mechanistic inference under sparse data conditions (3). We further investigate how neural architecture influences inverse problem robustness by comparing standard PINNs with PIKANs (66). Recent work suggests that architectures based on Kolmogorov-Arnold representations can improve expressivity and stability in nonlinear inverse problems (71; 18; 17). In addition to architectural design, we also examine implementation choices that affect inverse-problem stability, including collocation-point selection strategies, with additional experiments provided in the Appendix.

A novel aspect of our approach is the tight coupling between the DPD simulation and parameter inference in the ODE-based signaling model. Within this framework, PINNs estimate unknown model parameters, such as rate constants and binding affinities, by fitting the ODE system to partially observed trajectories while enforcing consistency with the governing dynamics. In the present study, this inverse analysis is primarily validated using synthetic observations together with DPD-derived trajectories and selected literature-based benchmarks. The PINNs framework learns the time evolution of system states (e.g., concentrations of signaling molecules) such that it not only matches available experimental measurements but also satisfies the ODE constraints (62; 38). This approach is particularly suitable in this context because experimental data on RBC phagocytosis signaling are sparse, often limited to a few time points from assays or microfluidic “spleen-on-a-chip” experiments (50; 90). The DPD simulations further enhance parameter inference by providing spatial–temporal constraints grounded in first-principles physics (37). For example, the DPD model can resolve CD47-SIRP*α* complex formation as a function of cell-cell distance and time, which can then inform ODE model initial conditions or serve as validation inputs (36; 46).

Using this hybrid modeling approach, we investigated how disease-associated perturbations in RBC surface signaling can influence macrophage phagocytosis. Rather than constructing patient-specific disease models, we considered representative perturbations motivated by SCD and GD that converge on a common functional consequence: attenuation of the CD47-SIRP*α* signaling complex *X*_3_. In this setting, the coupled DPD-ODE framework links experimentally motivated changes such as CD47 downregulation in SCD and enhanced opsonization in GD to reduced inhibitory signaling and altered downstream SHP1-mediated responses (22; 21; 28). These studies show how DPD-derived intermediate observables can be incorporated into predictive signaling dynamics, while the inverse analyses demonstrate that inclusion of *X*_3_ substantially improves trajectory prediction and parameter recovery (50).

In summary, we developed an integrated multiscale computational framework that combines molecular signaling with biophysical cell interactions to investigate RBC phagocytosis in both healthy and disease conditions. The approach couples DPD simulations of mesoscale membrane dynamics (25) with a mechanistic ODE model of the CD47-SIRP*α* signaling pathway, enabling explicit representation of antibody diffusion, membrane binding dynamics, and receptor activation during macrophage-RBC contact. The DPD simulation environment is physically calibrated by mapping simulation units to experimentally reported diffusion coefficients and validating receptor-ligand interaction dynamics, ensuring quantitative biological realism. To infer unknown kinetic parameters under sparse experimental observations, we incorporate an identifiability-aware inference framework that combines structural and practical identifiability analysis with PINNs (62), and we further examine the influence of neural architecture by comparing standard PINNs with PIKANs. Together, this integrated simulation–inference framework demonstrates how incorporating DPD-derived receptor activation trajectories can improve parameter recovery and reduce uncertainty in high-dimensional signaling models governing RBC phagocytosis. The framework further provides mechanistic insight into how disease-motivated perturbations of the CD47-SIRP*α*-SHP1 axis reshape inhibitory signaling and establishes a computational basis for exploring potential therapeutic modulation of this pathway (60).

## Materials and Methods

We developed an integrated multiscale modeling and inference framework combining a signaling ODE model, DPD simulations, identifiability analysis, and physics-informed neural networks. The DPD method was used to represent mesoscale membrane interactions and antibody–receptor binding dynamics, while the ODE model describes intracellular signaling pathways governing macrophage-mediated erythrophagocytosis. Identifiability analysis was performed to determine which kinetic parameters can be reliably inferred from available observables, and physics-informed neural networks were employed to estimate unknown parameters while enforcing consistency with the governing dynamics. To represent the biomechanics of blood cells and membrane interactions, we employ the DPD method, which has been widely used for mesoscopic simulations of blood cells and soft matter (42; 43; 50; 48; 11; 20; 77). In this approach, particles interact through pairwise forces that represent the chemical and physical interactions between different components. Details of the DPD formulation and parameter values used in this work are provided in the Supplementary Information.

### Signaling ODE Model

To describe intracellular signaling events governing erythrophagocytosis, we adopted and extended the systems biology framework introduced in (90), which links receptor-level interactions with downstream cytoskeletal regulation. The model incorporates both inhibitory and pro-phagocytic signaling pathways involved in macrophage-mediated clearance of red blood cells, including the CD47–SIRP*α* inhibitory pathway, SHP1 activation, Fc receptor signaling, phosphatidylserine receptor engagement, and myosin IIa dynamics. The signaling network is formulated as a system of 15 coupled nonlinear ordinary differential equations describing the temporal evolution of receptor complexes and intracellular signaling states. For clarity, the governing equations are summarized in Table S1. In the signaling model, *X*_3_ represents the activated CD47– SIRP*α* receptor complex that mediates inhibitory signaling. In the DPD simulations, receptor occupancy is used as a mechanistic proxy for this activation state, under the assumption that the fraction of CD47–SIRP*α* binding events correlates with the activation level of the inhibitory signaling pathway. State variables *X*_1_–*X*_15_ represent receptor states and intracellular signaling intermediates summarized in Table S3, while parameter definitions and nominal values are provided in Table S2. This nonlinear dynamical system captures the coupled receptor binding, activation kinetics, and intracellular feedback mechanisms that regulate macrophage activation and RBC engulfment dynamics. A detailed list of DPD equations and parameters—including interaction potentials, diffusion time scales, particle densities, and receptor placement methods—is provided in the Supplementary Information (SI). All simulations were performed using an extended version of the code developed in LAMMPS. Each simulation takes about 4,000,000 time steps to 8,000,000 time steps. A typical simulation requires 1200 CPU core hours to 2400 CPU core hours by using the computational resources (Intel Xeon E5-2670 2.6 GHz 24-core processors) at the Center for Computation and Visualization at Brown University.

### Structural Identifiability Analysis

We performed a *structural identifiability analysis* to determine which parameters in the signaling ODE model can be uniquely inferred from the available observables. Following the rational-form strategy of (16), the governing equations were reformulated to ensure rational dependence on the unknown parameters. In particular, the second and third equations contain the term

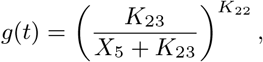

where *K*_22_ appears as an exponent. Because the algorithms implemented in `StructuralIdentifiability.jl` require rational parameter dependence, this transcendental form cannot be analyzed directly. Therefore, *K*_22_ was fixed to its experimentally determined value (*K*_22_ = 3) and treated as known during the identifiability analysis. Two analysis scenarios were considered: (i) all kinetic and affinity parameters treated as unknown except *K*_22_, and (ii) a reduced set in which experimentally measurable parameters were fixed. Parameters were classified as *globally identifiable, locally identifiable, nonidentifiable*, or *fixed* based on the symbolic analysis results.

### Practical Identifiability Analysis

Practical identifiability was evaluated to determine the recoverability of model parameters under finite and noisy measurements (64; 79). To mimic experimental variability, multiplicative noise was introduced into observable states at three levels (1%, 5%, and 10%) using independent uniform perturbations. No artificial noise was added to the experimentally measured CD47–SIRP*α* activation state (*X*_3_). Based on the results of the structural identifiability analysis, a subset of identifiable kinetic parameters was estimated using nonlinear least-squares optimization in log-parameter space, which ensures parameter positivity and improves numerical stability. The objective function was defined as a weighted sum of squared residuals between model predictions and observations, normalized by the magnitude of each state variable.

Local practical identifiability was first assessed using the Fisher Information Matrix (FIM), which was computed from the sensitivity matrix of model outputs with respect to the unknown parameters (64; 79). The resulting matrix was used to evaluate parameter variances, parameter correlations, and the conditioning of the estimation problem.

To complement the local FIM-based analysis, profile likelihood analysis was performed to examine parameter identifiability more globally. In this procedure, each parameter was fixed sequentially across a range of values in log-space while the remaining parameters were re-optimized (64). The increase in the objective function was then compared with the *χ*^2^ threshold corresponding to a 95% confidence level. Parameters exhibiting finite profile widths were considered practically identifiable, whereas flat or unbounded profiles indicated non-identifiability. Together, the FIM and profile likelihood analyses provide complementary local and global perspectives on parameter identifiability in the nonlinear ODE system.

### Physics-Informed Networks (PINs)

To estimate unknown kinetic parameters while enforcing mechanistic consistency with the governing signaling dynamics, we employed Physics-Informed Networks. In this framework, the state trajectories are approximated by a trainable function 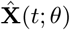, and the ODE system is embedded directly into the training objective through automatic differentiation.

As illustrated in Figure 3, the network receives time *t* as input and outputs approximations of the 15 signaling states. Temporal derivatives required for enforcing the ODE constraints are computed via automatic differentiation, allowing evaluation of the residual of the governing system at selected collocation points. Additional experiments evaluating the effect of collocation point selection (fixed grids versus adaptive resampling) on parameter recovery and training stability are presented in Appendix.

**Figure 1.**
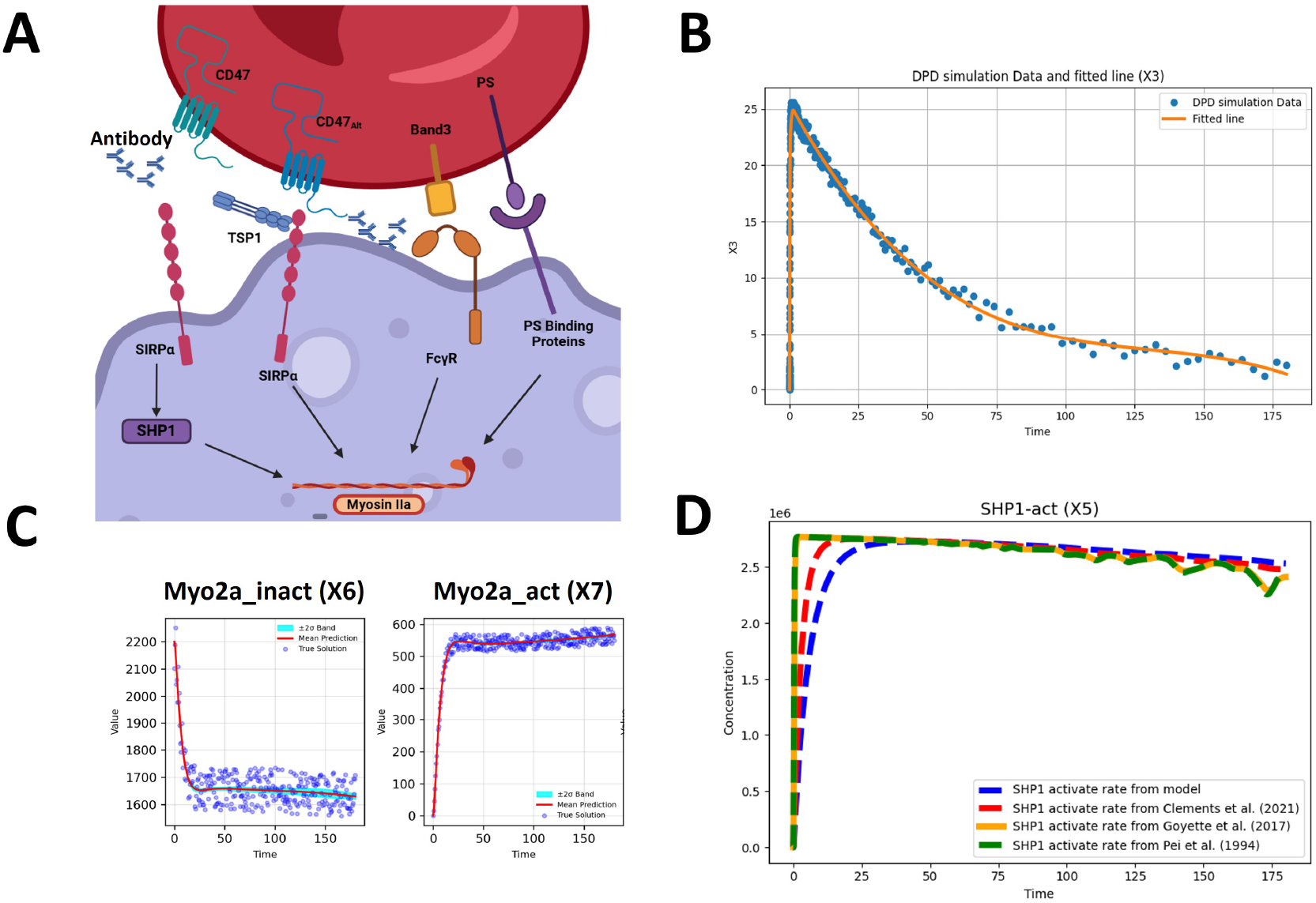
Integrated modeling and data analysis of RBC–macrophage signaling and SHP1 dynamics. (A) Schematic illustration of the recognition and signaling pathways regulating RBC clearance. Interactions between CD47-SIRP*α*, phosphatidylserine (PS)-binding proteins, Fc*γ*R, and TSP1 mediate phagocytosis signaling. Downstream, SHP1 activation and myosin IIa regulation contribute to engulfment. (B) Dissipative particle dynamics simulation data for binding dynamics (X3) compared with fitted line. (C) Physics-informed neural network inference of Myosin IIa dynamics. Left: inactivated Myo2a (Myo2a inact); Right: activated Myo2a (Myo2a act). Mean prediction (red) with ±2*σ* uncertainty band (cyan) is compared with noisy synthetic data (blue dots) and the true solution (black). (D) Comparison of SHP1 activation (X5) kinetics. Model-derived activation rate (blue) is benchmarked against reported values from Clements *et al*. (2021, red), Goyette *et al*. (2017, orange), and Pei *et al*. (1994, green) (15; 29; 57).

**Figure 2.**
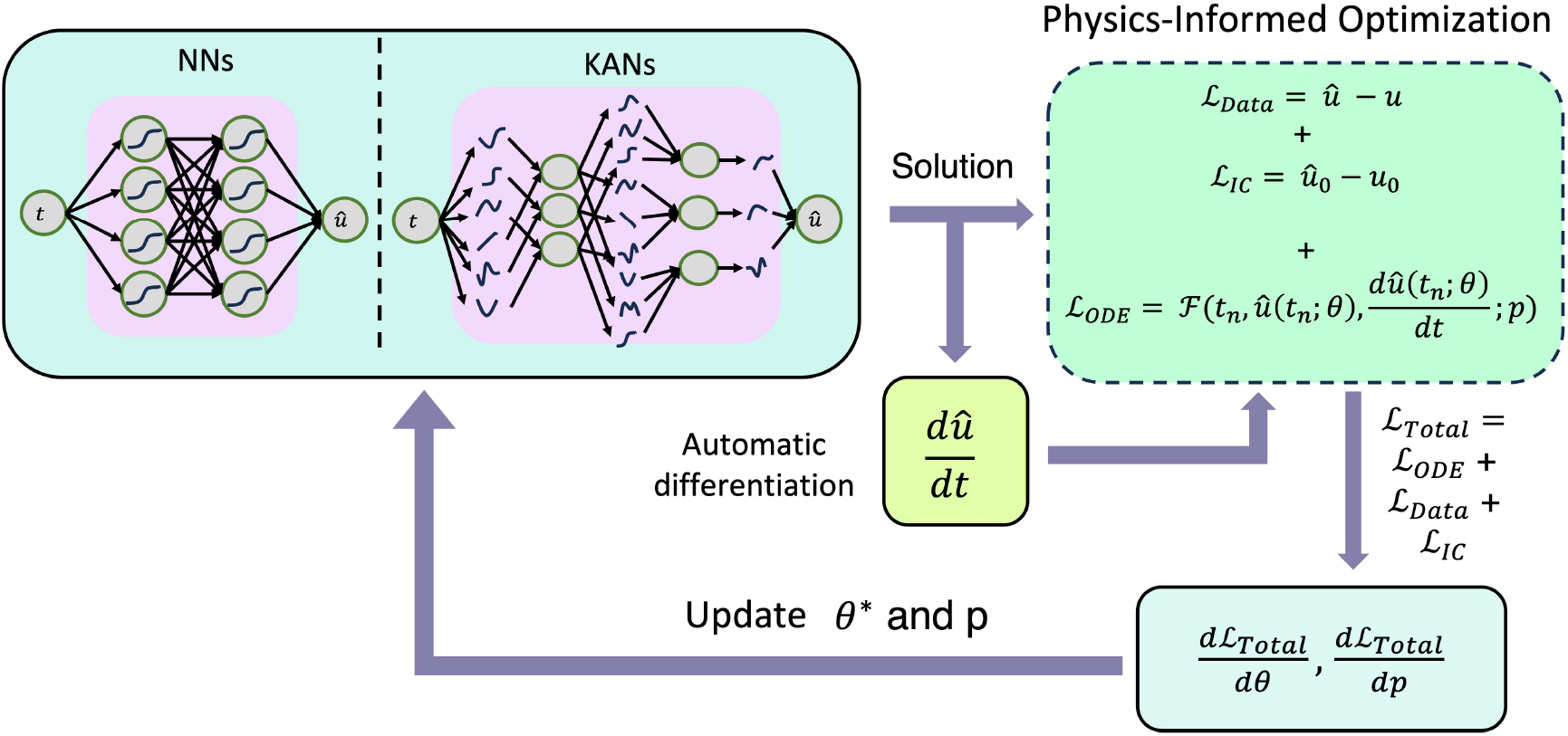
Schematic of the physics-informed networks (PINs) training framework. The network block represents the choice of network architecture used to approximate the system dynamics. Depending on the implementation, this block can employ either neural networks (NNs) or Kolmogorov-Arnold networks (KANs). The selected architecture is then trained within the physics-informed learning framework, where data-driven loss terms are combined with governing equation constraints to guide the optimization. *θ* denotes the trainable parameters of the network, and *p* represents the unknown parameters of the governing mathematical model.

**Figure 3.**
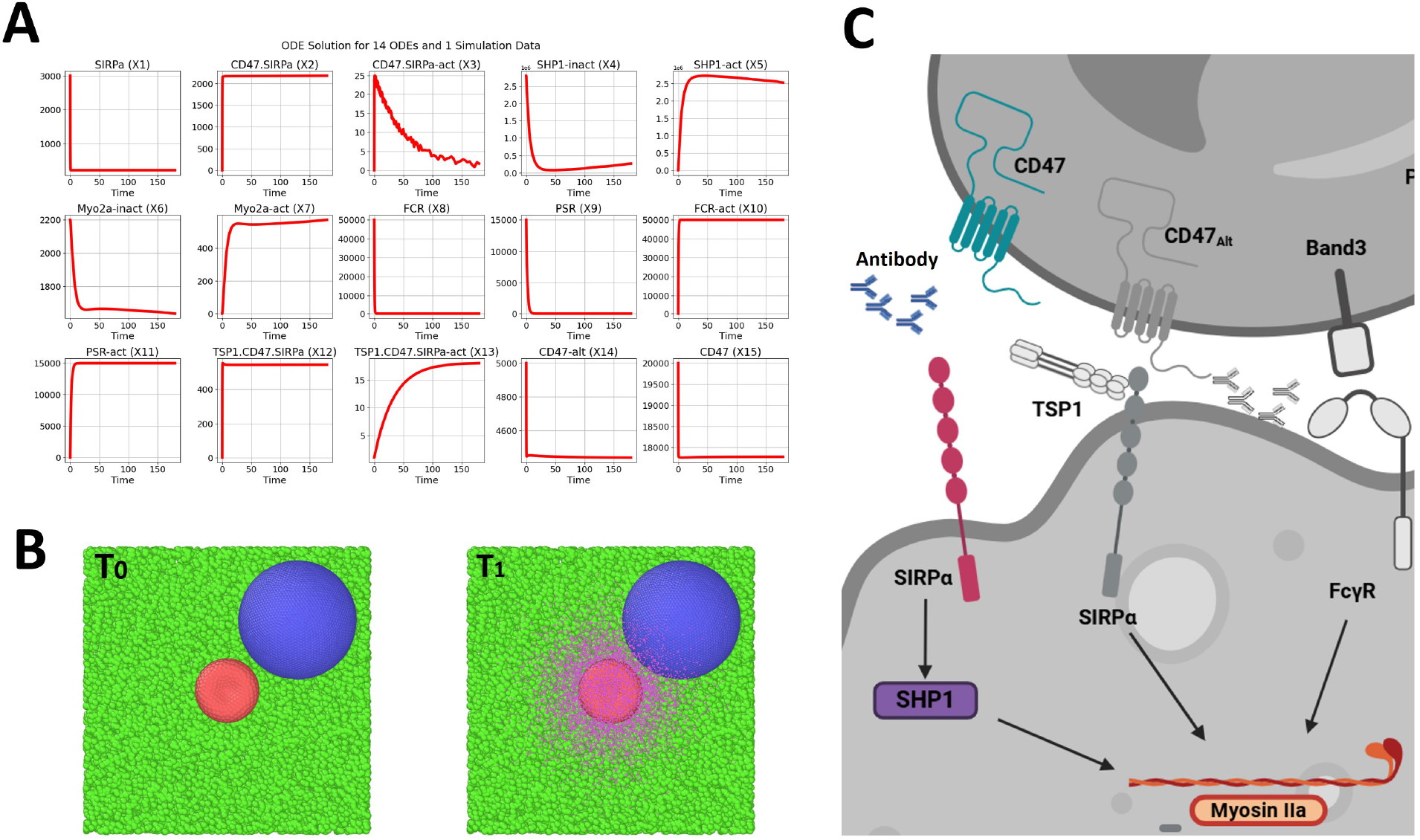
Integrated ODE-based signaling dynamics, particle simulations, and mechanistic signaling pathways. (A) Time-course solutions of a 15-variable ODE system modeling the dynamics of RBC-macrophage signaling. X3 (CD47-SIRP*α* complex) includes interpolated experimental data for parameter fitting. (B) DPD simulation snapshots showing RBC (red) and macrophage (blue) membrane interactions within a green fluid particle environment at time points *t* = 0 s and *t* = 1 s. Diffusing signaling molecules and particle-level interactions are visualized. (C) Mechanistic diagram of the CD47-SIRP*α*-SHP1 signaling axis, with inactive molecules grayed out to indicate early time points prior to engagement.

The total loss function is defined as

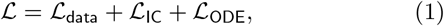

where ℒ_data_ penalizes the mismatch between model predictions and measured observations, ℒ_IC_ enforces consistency with the initial conditions listed in Table S3, and ℒ_ODE_ enforces the residual of the governing ODE system evaluated at the collocation points. Within this framework, we compare two representation models for 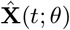: (i) a standard multilayer perceptron (MLP), and (ii) a Chebyshev-based Kolmogorov–Arnold Network with nested tanh nonlinearities (tanh-cPIKANs), introduced and systematically studied in (18; 40), where it was shown to outperform the standard Chebyshev-based Kolmogorov–Arnold architecture, particularly in solving ODE-based inverse problems. Additional architectural details are provided in the Appendix.

### Inverse Problem Setup

Following the structural identifiability analysis, we formulated the inverse problem by restricting attention to a subset of parameters deemed estimable under the available measurement configuration. We assume that nine of the fifteen state variables are experimentally measurable. In addition, the time-resolved trajectory of *X*_3_ is obtained from the DPD simulations. Consequently, observational data are available for the following ten state variables:

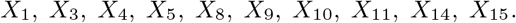

The parameters selected for estimation are

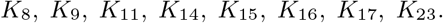

Because complete experimental measurements are not available for all observable states, synthetic data were generated by solving the forward problem. It enables controlled validation of identifiability and inference performance, which is not feasible with sparse experimental datasets. This procedure provided synthetic observations for the measurable states as well as ground-truth trajectories for the unobserved state variables to enable controlled validation of parameter recovery. The DPD-derived trajectory for *X*_3_ was incorporated into the governing equations, and the full coupled ODE system was integrated forward in time to produce dynamically consistent synthetic data for the remaining observable states. The resulting solution trajectories are shown in Figure 2A.

The parameter set is summarized in Table S2, which specifies the nominal values used for forward simulation, identifies the parameters selected for inverse estimation, and provides their corresponding search ranges. The initial conditions employed in the forward problem are listed in Table S3. During the inverse analysis, only the designated unknown parameters were estimated. For parameters with available prior information, predefined search intervals were imposed as indicated in Table S2. The remaining estimated parameters were allowed to vary freely during optimization, while all other parameters were fixed at their nominal values.

## Results

### Estimation and Validation of Diffusion Coefficients

In our multiscale framework, the diffusion and binding of antibody particles are simulated specifically during the phagocytosis. This localized process is initiated at *T* = 0, corresponding to the onset of sensing between the macrophage pseudopodia and the red blood cell (RBC) membrane. Figures 3B and 3C illustrate the receptor–ligand dynamics and representative RBC–macrophage configurations obtained from the DPD simulations. Rather than simulating the computationally expensive bulk diffusion of antibodies in the entire fluid domain, we initialize the therapeutic particles at the RBC boundary. This specific initial condition represents the localized high effective concentration of opsonizing antibodies confined within the narrow space immediately following *T* = 0. As these particles dynamically diffuse and bind to the macrophage surface receptors in the DPD simulation, they progressively occupy the available binding sites. This physical occupancy induces steric hindrance, preventing native CD47-SIRP*α* ligation. Consequently, the DPD-derived spatial receptor occupancy serves as a spatiotemporal boundary condition that mechanistically dictates the temporal evolution of the inhibitory signaling complex, *X*_3_(*t*), in our ordinary differential equation (ODE) system. By employing this multiscale coupling, our model bridges the microsecond-scale spatial constraints with the minute-scale intracellular signaling dynamics without relying on well-mixed assumptions. This approach inherently captures the mechanosignal coupling where cellular physical contact precedes and governs localized biochemical signaling (67).

To characterize the diffusive behavior captured by the DPD simulations, we analyzed particle trajectories, radial distributions, and mean squared displacement (MSD) statistics, as illustrated in Figure S1. Representative particle trajectories in Figure S1A exhibit typical three–dimensional random walk behavior. As diffusion progresses, particles spread radially from their initial location, leading to the isotropic density distribution shown in Figure S1B. The radial symmetry of this distribution confirms that the simulated transport is consistent with Brownian diffusion.A quantitative characterization of diffusion is obtained from the MSD curves shown in Figure S1C, which correspond to three representative particle sizes (10 nm, 12 nm, and 15 nm).

### Validation of Antibody Dynamics

The interaction between CD47 on target cells and SIRP*α* on macrophages provides a critical inhibitory “don’t eat me” signal that suppresses phagocytosis (80; 60). Engagement of CD47 with SIRP*α* activates inhibitory signaling in macrophages, preventing the clearance of self cells (14). In our multiscale signaling model, this inhibitory pathway is represented by the activation state of the CD47–SIRP*α* complex, denoted as *X*_3_. Direct observation of antibody diffusion and receptor binding at cell membranes remains experimentally challenging due to nanometer spatial and microsecond temporal scales (72). To investigate these processes and supply critical inputs to our ODE signaling model, we employed DPD simulations. To demonstrate the versatility of our framework, we investigate the disrupted phagocytic signaling observed in both GD and SCD. While GD and SCD have fundamentally different pathophysiological origins, both conditions are characterized by a compromised CD47–SIRP*α* inhibitory axis. In GD, experimental studies reveal a general decrease in membrane CD47 expression on red blood cells (RBCs) by 1% to 26% compared to normal cells (9). Concurrently, GD RBCs face heightened opsonization due to elevated autoantibodies (IgG1, IgG2, IgG3) (56), with patient samples showing up to a twofold increase in antibody binding despite similarities in baseline CD47 levels (21). Conversely, in SCD, the disrupted signaling is primarily driven by an inherent reduction of CD47 expression on the RBC surface, while macrophage SIRP*α* expression remains normal (22).

Our primary objective is to establish a unified computational framework that effectively connects DPD-derived observables to the ODE signaling model. To demonstrate the biological utility of this approach, we utilized DPD simulations to capture a critical functional consequence common to both GD and SCD: the attenuation of the *X*_3_ signaling complex. Figure S2 summarizes these results across temporal, spatial, and signaling dimensions. By utilizing a representative DPD simulation to model the lowered *X*_3_ activation, we demonstrate how our coupled DPD-ODE framework can successfully incorporate experimental disease markers—whether originating from autoantibody opsonization in GD or CD47 downregulation in SCD—into predictive signaling dynamics for macrophage phagocytosis.

### Validation of Downstream Signaling Dynamics through SHP1 Activation and Pathway Crosstalk

To further validate our multiscale signaling framework, we examined the activation dynamics of SHP1, a key downstream effector of CD47–SIRP*α* interactions, and compared simulation outputs with experimental measurements. Figure 4 summarizes these results through both quantitative ODE solutions and a schematic overview of the signaling cascade.

**Figure 4.**
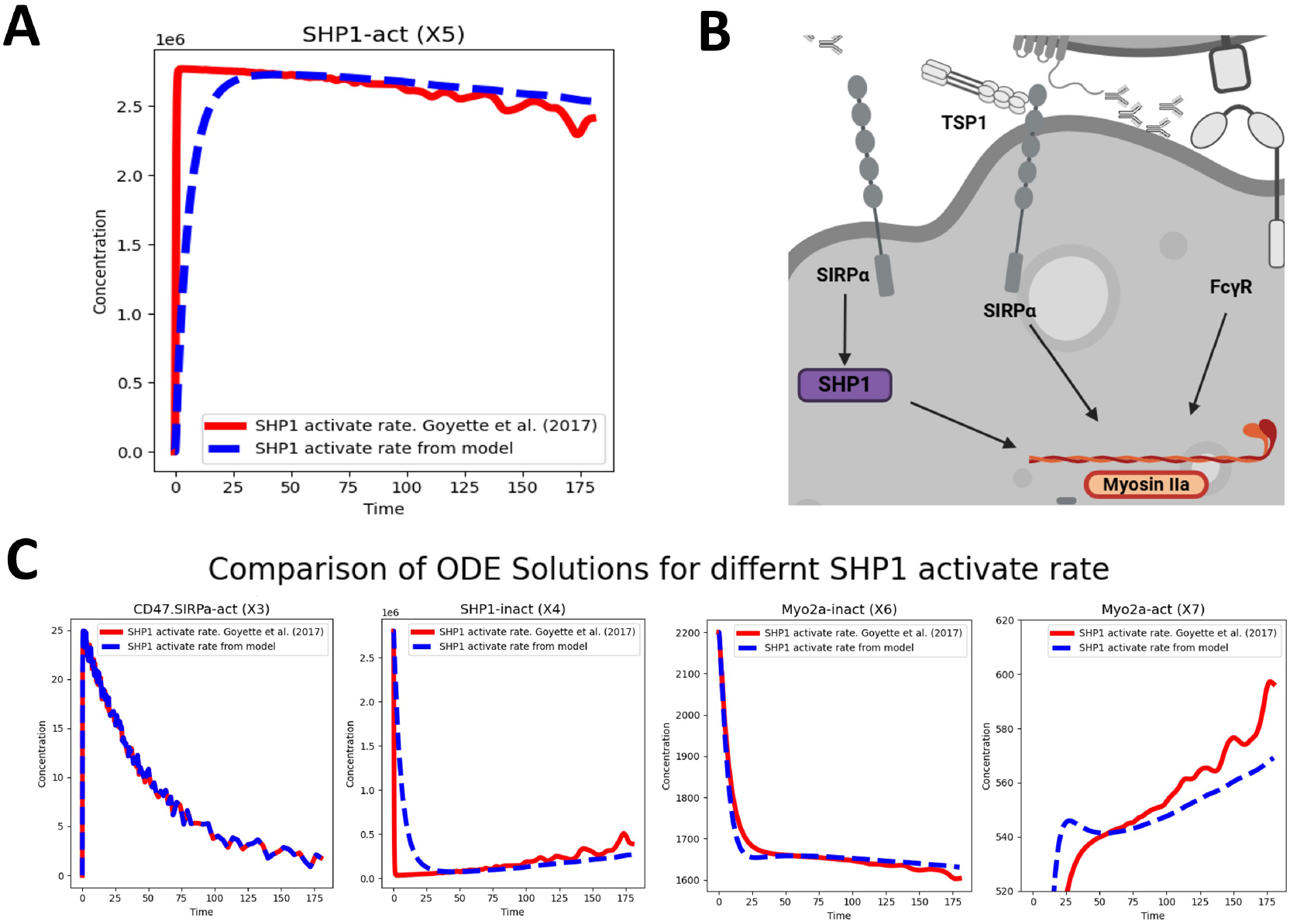
Effect of SHP1 activation rate on RBC clearance signaling dynamics. (A) Comparison of SHP1 activation (X5) based on the model (blue) and the experimentally reported rate from Goyette et al. (2017, red). (B) Schematic diagram of the SHP1-mediated signaling network downstream of SIRP*α*, highlighting the effect of SHP1 activation rate on Myosin IIa activation. (C) Comparison of ODE simulation results for key variables (X3: CD47-SIRP*α* complex activation, X4: SHP1 inactivation, X6: Myo2a inactivation, X7: Myo2a activation) for different SHP1 activation rates. In each case, the model-derived solution (blue) is benchmarked against the rate from Goyette et al. (2017, red), showing how SHP1 kinetics influence molecular responses.

Figure 4A shows the time evolution of activated SHP1 (X5) in response to receptor engagement. The model predicts a rapid increase in SHP1 activation, reaching a steady state within the first 50 time units. This trajectory closely follows experimental measurements reported by Goyette et al. (2017), with the model reproducing both the sharp onset of activation and the subsequent plateau. The agreement indicates that the estimated SHP1 activation rate in our framework captures the experimentally observed dynamics. The pathway schematic in Figure 4B highlights how SIRP*α* engagement recruits and activates SHP1, which in turn modulates downstream effectors including Fc*γ*R and Myosin IIa. This cascade forms the inhibitory arm of the phagocytic checkpoint, integrating antibody blockade, receptor engagement, and cytoskeletal remodeling. To probe the robustness of this pathway representation, we compared ODE solutions for multiple downstream species under different SHP1 activation rates. Figure 4C shows CD47-SIRP*α*-act (X3), SHP1-inact (X4), Myo2a-inact (X6), and Myo2a-act (X7). In each case, the red curves represent experimental activation dynamics from Goyette et al. (29), while the blue curves show simulation results. Across all readouts, the model reproduces the qualitative temporal patterns: rapid depletion of inactive pools (X4, X6), concurrent accumulation of active forms (X5, X7), and sustained reduction of receptor–ligand complexes (X3). Quantitative discrepancies at later times likely reflect simplifications in our current parametrization of SHP1 turnover and cytoskeletal feedback, which can be addressed in future iterations by incorporating experimentally measured dephosphorylation rates and actin remodeling constraints. Overall, these results demonstrate that the model captures the essential features of SHP1-mediated inhibitory signaling. The agreement with experimental datasets, combined with the mechanistic pathway representation, validates the use of our ODE framework to study antibody-mediated modulation of SIRP*α* signaling and its downstream cytoskeletal consequences.

### Structural and Practical Identifiability Analysis

Structural identifiability analysis was performed to determine which kinetic parameters can, in principle, be uniquely recovered from ideal noise-free measurements under the chosen observable configuration. The results under three modeling scenarios are summarized in Table S5. Importantly, this reduction step does not bias the inference process; rather, it ensures that only theoretically recoverable parameters are estimated from data. However, structural identifiability alone does not guarantee reliable parameter recovery in practice. Real experimental measurements are finite, noisy, and often incomplete. Even parameters that are structurally identifiable under ideal noise-free conditions may become weakly constrained when estimation is performed with limited data and measurement uncertainty. Therefore, we next assess practical identifiability, which quantifies the extent to which parameters can be estimated with acceptable uncertainty under realistic observational noise.

We evaluated practical identifiability of the selected kinetic parameters under three observational noise levels (1%, 5%, and 10%). Parameter estimation was performed using nonlinear least-squares optimization, followed by local curvature analysis via the FIM and global assessment through profile likelihood. Across all noise levels, the optimization converged to finite-cost solutions, although deviations from nominal parameter values increased as noise amplitude increased. The FIM condition numbers were on the order of 10^5^–10^6^, indicating substantial ill-conditioning and suggesting the presence of poorly constrained parameter combinations. The FIM correlation heatmaps (Figure 5) show a stable correlation structure across noise levels.

**Figure 5.**
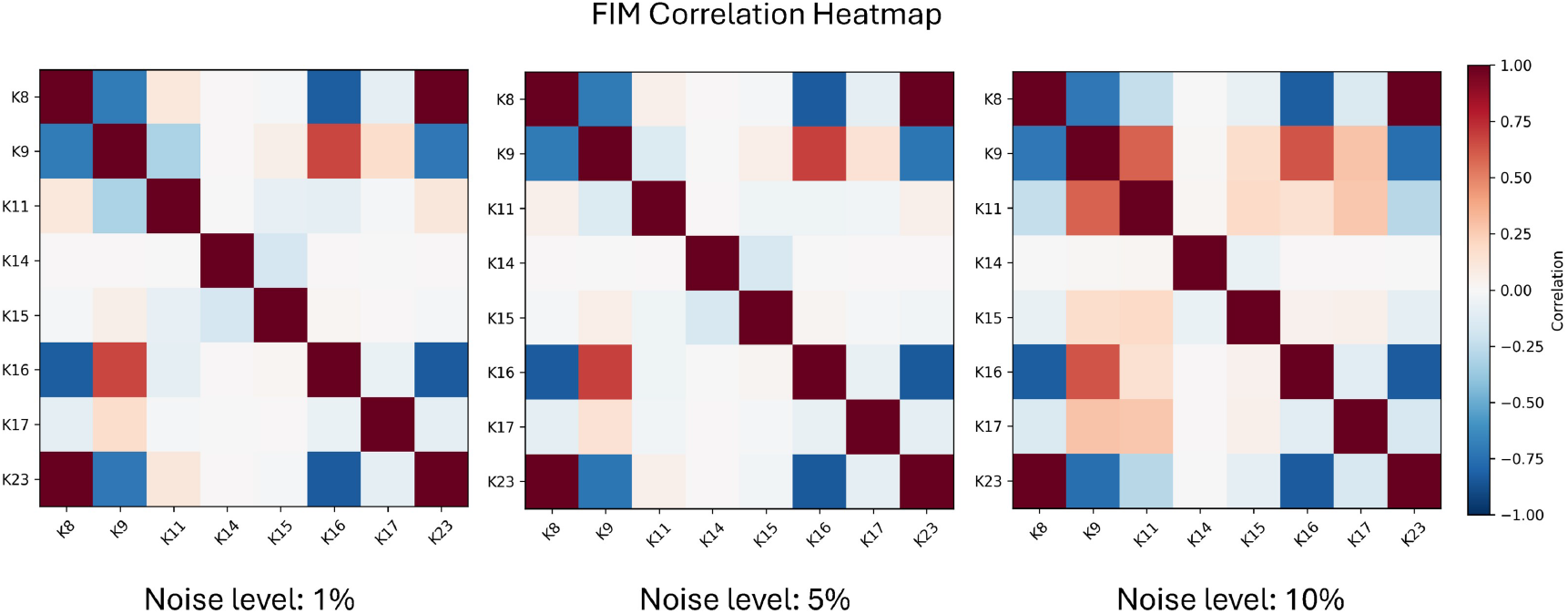
Fisher Information Matrix (FIM) correlation heatmaps for the selected structural identifiable kinetic parameters under 1%, 5%, and 10% observational noise levels (left to right). Each panel displays the normalized parameter correlation matrix computed at the optimal fit. Strong positive correlation is consistently observed between *K*_8_ and *K*_23_ and between *K*_9_ and *K*_16_, while *K*_8_ and *K*_16_ exhibit pronounced negative correlation. The overall correlation structure remains stable across noise levels, indicating intrinsic parameter coupling in the model rather than noise-induced artifacts.

The agreement between FIM-based curvature diagnostics and profile likelihood confirms the distinction between locally and globally constrained parameters. Table S6 summarizes the fitted parameter values, FIM-based standard deviations, profile likelihood outcomes, and agreement assessment across noise levels. Collectively, these results demonstrate that while several parameters remain practically identifiable under moderate noise, specific kinetic rates, particularly *K*_11_ and *K*_15_, are weakly informed by the available observables and are highly sensitive to measurement uncertainty.

### Effect of Search Space Constraints on Parameter Stability

The practical identifiability analysis revealed substantial ill-conditioning in the reduced parameter set, characterized by large FIM condition numbers (~ 10^5^–10^6^), strong parameter correlations (notably between *K*_8_ and *K*_23_), and weak curvature directions for *K*_11_. These findings indicate the presence of compensation effects and flat likelihood directions that can destabilize unconstrained optimization.

To mitigate variance inflation and compensation-induced parameter drift, we introduced search limits for parameters *K*_8_, *K*_11_, and *K*_23_, which showed the strongest instability in the practical identifiability analysis. All remaining parameters were allowed to vary freely within broad admissible ranges.

Table S7 compares parameter estimates obtained with and without search limits. Without constraints, significant bias and dispersion are observed for the red-highlighted parameters. In particular, *K*_8_ is overestimated (mean 7.16 × 10^−1^ vs. true 4.00 × 10^−1^) with substantial variability, and *K*_11_ exhibits nearly threefold overestimation and large standard deviation (7.02 × 10^−4^). Similarly, *K*_23_ drifts downward with inflated variance.

Imposing search limits substantially improves both accuracy and stability. For *K*_8_, the mean estimate shifts closer to the true value (3.83 × 10^−1^) and the standard deviation decreases by more than fourfold. The effect is even more pronounced for *K*_11_, whose standard deviation collapses from 7.02 × 10^−4^ to 9.00 × 10^−6^, indicating removal of a flat curvature direction. Likewise, the variance of *K*_23_ decreases by approximately 75%, and its mean estimate aligns closely with the nominal value.

Importantly, parameters that were already well-conditioned (e.g., *K*_9_, *K*_14_, *K*_16_, *K*_17_) remain stable under both settings, as we expected that the imposed limits do not artificially bias well-identified directions.

The results show that search space restriction, when guided by identifiability diagnostics, acts as a regularization mechanism that suppresses compensation effects and stabilizes inverse inference without compromising mechanistic interpretability.

### Impact of Incorporating *X*_3_ Simulation Results on Prediction and Parameter Inference

A key question in modeling complex biological signaling systems is whether incorporating additional intermediate-state observables can improve predictive performance. In our system, *X*_3_ corresponds to the activated CD47-SIRP*α* complex, a central hub in the inhibitory signaling cascade. Biologically, this state represents the intracellularly active form of the CD47–SIRP*α* immune checkpoint, which plays a crucial role in regulating phagocytosis of red blood cells by macrophages. Although such intermediate complexes are rarely measured directly in experiments due to technical limitations, they provide mechanistically rich information that could anchor the model and constrain parameter estimation. The simulation results enable controlled validation of identifiability and inference performance, which is not feasible with sparse experimental datasets. We therefore investigated how the inclusion of *X*_3_ as an observable affects the outcome of PINNs–based parameter inference. To quantify these effects, we designed an uncertainty quantification (UQ) study where each training scenario was repeated 10 times with different random seeds. This procedure captures stochastic variability from initialization and optimization, providing not only average predictions but also distributions of possible outcomes. Such UQ analyses, widely used in physics-informed machine learning frameworks (85; 49), offer a rigorous means of assessing both accuracy and robustness. To further stabilize PINNs training, we employed a smooth *q*-scheduling strategy that gradually increases the weight of the physics-based loss relative to the data loss during optimization; a detailed comparison of different *q*-scheduling strategies is provided in Figure S3 in the Supplementary Information.

Figures 6 and 7 summarize the results, obtained using the PINNs framework with 1000 residual points and 100,000 training epochs. Figure 6 shows predicted trajectories of all model species, with *X*_3_ highlighted as a representative readout. Without incorporating *X*_3_ simulation results (Figure 6A), trajectories diverge from the ground truth (red), but this discrepancy largely reflects parameter misestimation rather than dynamics alone. In particular, parameters such as *K*_8_, *K*_16_, and *K*_23_ exhibit strong bias, which propagates into large steady-state errors and inter-run variance reaching nearly 20%. Including *X*_3_ simulation results (Figure 6B) restores close agreement in trajectories, with deviations below 10% and markedly reduced variance, reflecting the improved parameter recovery.

**Figure 6.**
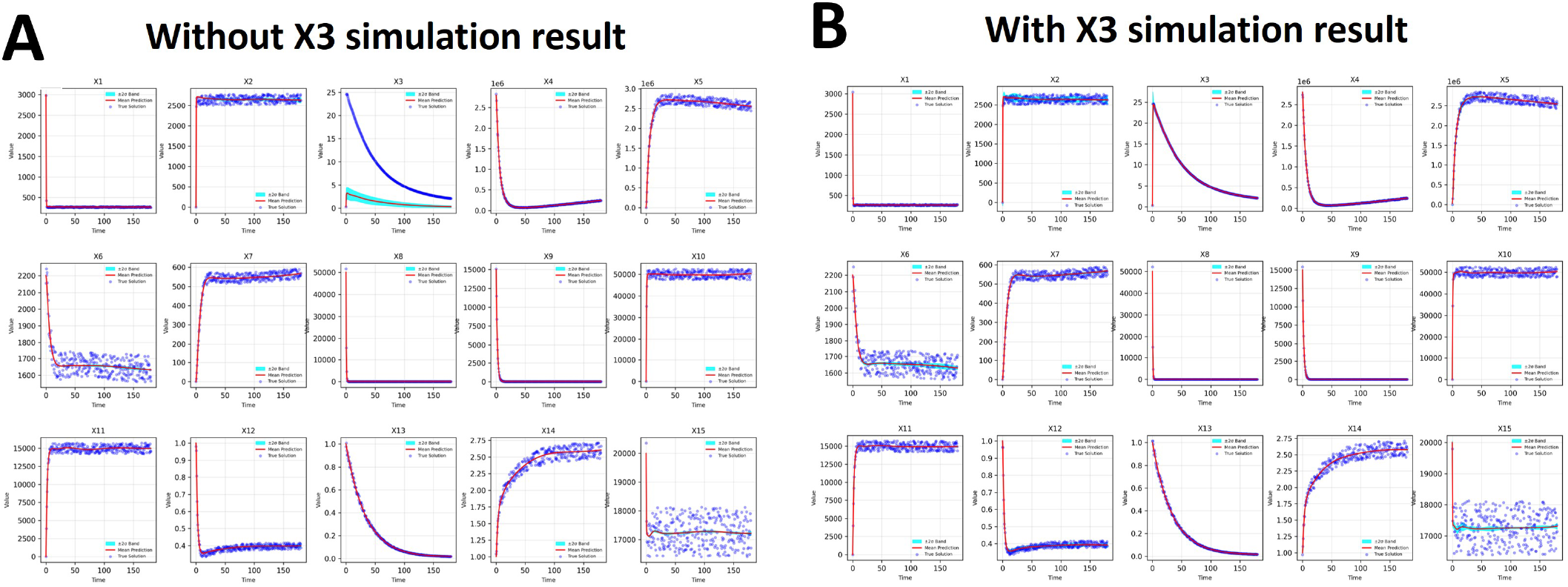
Impact of incorporating *X*_3_ simulation results on model prediction. (A) Model predictions (mean ± SD, red) without *X*_3_ simulation result, compared to ground truth (blue). (B) Model predictions with *X*_3_ simulation result, showing improved accuracy and reduced uncertainty across runs.

**Figure 7.**
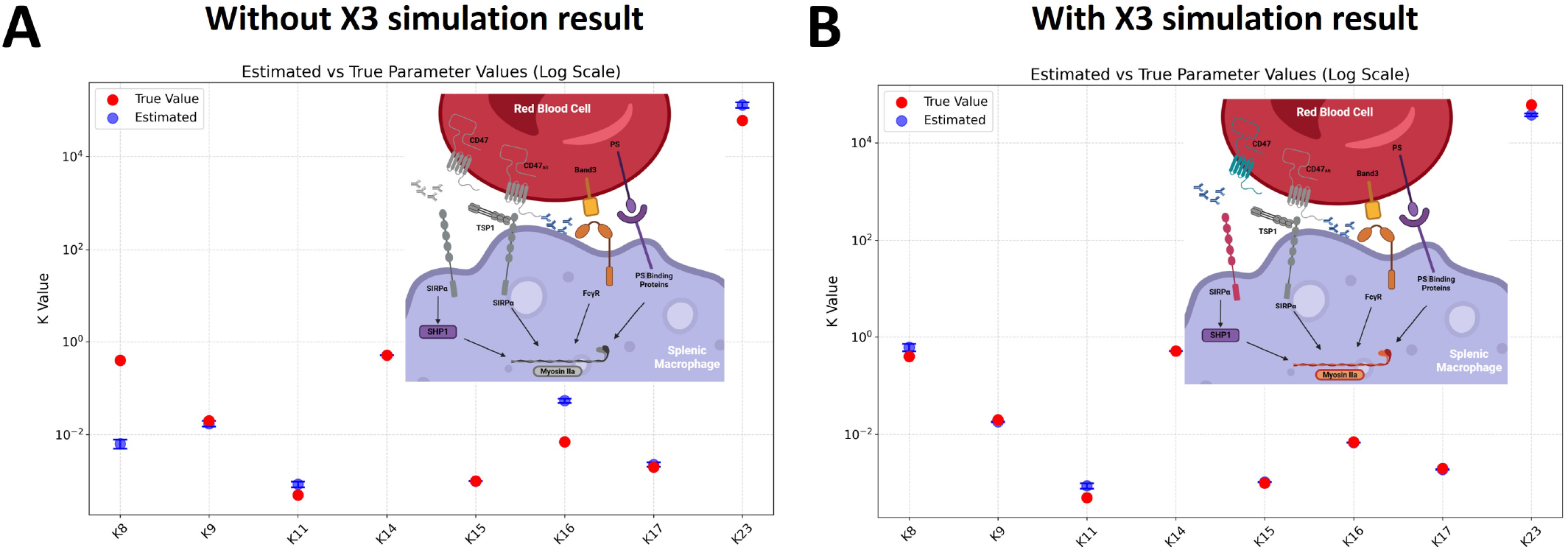
Impact of incorporating *X*_3_ simulation results on parameter inference. (A) Corresponding parameter estimates (blue) with standard deviation bars, plotted against true parameter values (red) on a logarithmic scale without *X*_3_ simulation result. (B) Parameter inference with *X*_3_ simulation result, showing reduced variance and improved agreement with ground truth, particularly for key parameters (*K*_8_, *K*_16_, and *K*_23_). Insets in (A, B) illustrate the biological context of the CD47-SIRP*α*-SHP1 pathway.

Figure 7 directly compares inferred kinetic parameters against their true values. Without *X*_3_ simulation results (Figure 7A), estimates for several parameters—notably *K*_8_, *K*_16_, and *K*_23_—exhibit large bias and variance. For example, *K*_8_ is underestimated by nearly an order of magnitude, with a standard deviation spanning 1.4 log-units, while *K*_23_ estimates show variance exceeding 60% relative to the mean. Incorporating *X*_3_ simulation results (Figure 7B) markedly improves recovery: the bias in *K*_8_ falls below 15%, variance in *K*_16_ decreases by 95%, and variance in *K*_23_ shrinks by over 70%. Overall, parameter estimates cluster tightly around ground truth values, confirming improved identifiability (76) and robustness of the inverse modeling process (62). Thus, the inclusion of even a single mechanistic observable propagates constraints throughout the model, tightening the solution space and stabilizing predictions of unmeasured species. This view makes clear that the trajectory deviations in Figure 6 are driven by biased parameter estimates.

Taken together, these results demonstrate that *X*_3_, although representing only a single intermediate species, has an outsized impact on both trajectory prediction and parameter inference. Its inclusion reduces error across the state space, dampens variability between training runs, and dramatically tightens posterior distributions of sensitive kinetic parameters. From a biological perspective, this suggests that targeted measurement of mechanistically central but experimentally challenging observables—particularly those associated with immune checkpoint nodes such as CD47–SIRP*α*—can substantially improve model-based inference. From a methodological perspective, it illustrates how PINNs-based approaches benefit from judiciously selected constraints: by anchoring the network to intermediate signaling states, the model is better able to recover hidden dynamics and identify parameter sets consistent with experimental reality. Overall, the study highlights that combining UQ with targeted observables provides a powerful strategy to achieve both predictive accuracy and reliable mechanistic interpretation in complex signaling models. This further suggests that, in experimental design, obtaining even a small number of strategically chosen measurements at key mechanistic nodes can dramatically increase the reliability of data-driven modeling.

### Robustness to Observation Noise

To assess the robustness of parameter inference under noisy conditions, we compared the performance of PINNs and Tanh-cPIKANs at different levels of additive observational noise: 0%, 5%, and 10%. For both methods, we used 1000 fixed collocation points uniformly distributed over the time interval *t* ∈ [0, 180].

To simulate a more realistic scenario, we perturbed synthetic measurement points using noise drawn from a uniform distribution. Specifically, we applied noise according to the following rule:

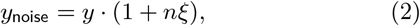

where *y* is the true model output, *n* is the noise level (e.g., 0.00, 0.05, 0.10), and *ξ* ~ 𝒰 (−1, 1) is a random variable sampled from a uniform distribution. This formulation ensures that the perturbed signals remain non-negative, which is critical when the underlying quantities are biologically constrained to be positive and may approach values close to zero.

Quantitative comparisons are presented in Table S8. At zero noise, the PINNs model generally achieves lower relative errors across most parameters. For instance, PINNs estimates *K*_11_ and *K*_15_ with errors of 0.76% and 0.18%, respectively, compared to 12.50% and 5.17% for tanh-cPIKANs. Similarly, PINNs shows lower errors for *K*_9_, *K*_17_, and *K*_23_ in the noise-free case. However, as observational noise increases, the relative performance of the two methods shifts. At 10% noise, tanh-cPIKANs show improved stability in several parameters. For example, the error in estimating *K*_9_ increases more sharply in PINNs (from 1.86% to 7.00%) than in tanh-cPIKANs (from 2.42% to 3.09%). Additionally, parameters like *K*_14_ and *K*_16_ remain within a tight error band (¡2.2%) for tanh-cPIKANs across all noise levels, whereas their PINNs counterparts exhibit more variability. It is worth noting that *K*_11_ remains better estimated by PINNs across all noise levels, with errors staying below 2.5%, while tanh-cPIKANs yield higher errors around 13%. Conversely, for some parameters such as *K*_8_ and *K*_17_, the two methods perform comparably or trade off depending on the noise level. Overall, while PINNs tend to outperform in low-noise settings, tanh-cPIKANs exhibit greater robustness and consistency under increasing noise—an important property when dealing with real-world data containing measurement uncertainty.

## Discussion and Summary

In this study, we show that incorporating physically derived intermediate observables (X3) fundamentally improves parameter identifiability and predictive accuracy in signaling models.

The integration of dissipative particle dynamics (DPD) simulations with PINNs addresses a fundamental challenge in systems biology: bridging the gap between mesoscale physical interactions and intracellular signaling dynamics. Traditional ordinary differential equation (ODE) models are effective in capturing average biochemical kinetics but often lack the spatial and stochastic resolution required to describe cell-cell interactions at the macrophage-RBC interface. By incorporating DPD-derived trajectories of receptor binding and molecular diffusion, our framework introduces physically grounded constraints into the signaling model, enabling a more realistic representation of how membrane mechanics and molecular transport influence signaling activation. In particular, the DPD-generated trajectory of the CD47-SIRP*α* complex activation state *X*_3_ provides an intermediate mechanistic observable that anchors the inverse problem and substantially improves parameter inference.

A key mechanistic insight emerging from this study is the central regulatory role of the CD47-SIRP*α*-SHP1 signaling axis in determining macrophage responses to red blood cells. The simulations demonstrate that small perturbations in the activation dynamics of SHP1 (*X*_5_) can propagate through the signaling cascade to significantly influence myosin IIa activation and the formation of the phagocytic cup. Because SHP1 acts as a phosphatase that counterbalances pro-phagocytic signaling pathways, its activation effectively determines whether macrophages remain in an inhibitory state or proceed toward engulfment. The identifiability results further reinforce this interpretation, showing that parameters governing SHP1 activation and CD47-SIRP*α* signaling are among the most influential and recoverable in the system.

The multiscale framework also provides insight into disease-specific mechanisms underlying pathological RBC clearance. In SCD, the inherent reduction of membrane CD47 expression directly weakens the inhibitory checkpoint signal. Within our multiscale model, this is captured through the attenuated activation of the CD47–SIRP*α* complex (*X*_3_). This weakened signal manifests as insufficient recruitment of SHP1 to suppress downstream cytoskeletal activation, allowing macrophages to initiate phagocytosis more readily—a result that aligns with the accelerated clearance of SCD RBCs in splenic microenvironments. Conversely, in GD, the compromised checkpoint originates from a combination of decreased CD47 expression and heightened opsonization by IgG autoantibodies. Despite these differing pathophysiological origins, our coupled DPD-ODE framework demonstrates how both conditions converge on a shared functional consequence: the reduction in *X*_3_ activation. By explicitly linking these disease-specific extracellular markers to the partial loss of the “don’t eat me” signal, the model provides a mechanistic explanation for how disrupted receptor engagements shift the intracellular signaling balance toward RBC engulfment. The DPD simulations indicate that these mechanical alterations modify receptor spatial distribution and effective binding affinity, which in turn shifts the signaling balance toward macrophage activation. These findings highlight the importance of considering both biochemical signaling and physical cell properties when investigating erythrophagocytosis. The simulations establish a quantitative framework for analyzing diffusion and localized binding dynamics. The results are consistent with fluorescence microscopy observations, where receptor–ligand complexes organize into microdomains with elevated local densities (74). While these results support the biological relevance of the model, it is important to note that the current framework does not include an explicit cytoskeletal scaffold. Such structures are known to restrict receptor diffusion and organize membrane domains in red blood cells and neurons (44; 45; 89; 10; 13), and their absence may contribute to quantitative differences between simulation and experiment. Nevertheless, by utilizing a representative DPD simulation to model the lowered *X*_3_ activation, we demonstrate how our coupled DPD-ODE framework can successfully incorporate experimental disease markers—whether originating from autoantibody opsonization in GD or CD47 downregulation in SCD—into predictive signaling dynamics for macrophage phagocytosis.

Beyond SCD and GD, the proposed multiscale framework provides a generalizable paradigm for understanding how mechanical and biochemical cues jointly regulate RBC clearance across diverse pathological conditions. By decoupling membrane mechanics from intracellular signaling while preserving their functional coupling through receptor-level interactions, the model enables systematic exploration of how distinct disease-specific perturbations converge on common immune checkpoint pathways. This generality is particularly relevant for infectious diseases such as malaria, where *Plasmodium*-infected RBCs exhibit both increased membrane rigidity and altered immune signaling. Previous studies suggest that parasites can exploit the CD47–SIRP*α* axis to evade macrophage clearance by enhancing “don’t eat me” signaling (6). Within our framework, such effects can be interpreted as a shift in both mechanical constraints and effective inhibitory signaling capacity, allowing the model to predict how competing influences—rigidity-induced retention versus signaling-mediated immune evasion—govern clearance outcomes. Similarly, in metabolic disorders such as diabetes, chronic oxidative stress leads to progressive stiffening of RBC membranes and a concomitant reduction in CD47 expression due to enhanced vesiculation (69). These coupled alterations naturally map onto our framework as simultaneous changes in DPD-resolved mechanical properties and baseline inhibitory signaling levels. Importantly, this unified representation suggests that seemingly distinct pathological processes may share a common mechanistic pathway: the disruption of spatially regulated CD47–SIRP*α* signaling at the macrophage interface. Taken together, these examples highlight that the predictive capability of the framework does not rely on disease-specific reparameterization of the entire model, but rather on targeted modulation of physically interpretable variables. This positions the model as a versatile platform for investigating how diverse biochemical and biomechanical perturbations reshape immune recognition and phagocytic decision-making.

From a methodological perspective, the comparison between standard PINNs and PIKANs reveals complementary advantages. While PINNs tend to achieve lower errors under low-noise conditions, PIKANs exhibit greater robustness to observational noise and high-frequency fluctuations arising from particle-based simulations. This robustness likely arises from the localized functional representation enabled by the Kolmogorov-Arnold architecture, which can capture localized nonlinear features more effectively than global multilayer perceptrons. As biological measurements often contain substantial noise and variability, these results suggest that hybrid architectures such as PIKANs may offer advantages in practical applications of physics-informed machine learning to biological systems.

Overall, this study demonstrates the feasibility and utility of combining particle-based biophysical simulations, mechanistic signaling models, and physics-informed machine learning into a unified multiscale framework. By explicitly linking receptor-level interactions, intracellular signaling dynamics, and parameter inference, the model provides a mechanistic explanation for dysregulated RBC clearance in SCD and GD. More broadly, the framework establishes a computational platform capable of exploring therapeutic interventions targeting macrophage immune checkpoints and offers a generalizable approach for studying multiscale immune signaling processes.

Despite the insights provided by the proposed framework, several limitations should be noted. The signaling network is modeled using deterministic ordinary differential equations that assume spatial homogeneity inside the macrophage, which may overlook spatial heterogeneity and stochastic effects in intracellular signaling. Parameter inference is mainly based on synthetic datasets with limited experimental validation, and some parameters were fixed to address structural non-identifiability, which may influence model predictions under different biological conditions. In addition, the DPD simulations use a coarse-grained representation of membranes and antibodies, which improves computational efficiency but simplifies molecular details such as cytoskeletal structures and receptor clustering. These assumptions were necessary to ensure computational tractability while retaining the dominant mechanisms governing macrophage-RBC signaling.

Future work will focus on improving biological realism and predictive capability by integrating additional experimental data, such as microfluidic spleen-on-a-chip assays or imaging of macrophage-RBC interactions, to better calibrate and validate the model. Extending the framework to include spatially resolved intracellular signaling, for example through reaction-diffusion or agent-based models, could capture spatial heterogeneity during phagocytic cup formation. Further development of physics-informed machine learning approaches, including uncertainty-aware methods, may also enhance parameter inference and predictive reliability. Finally, the framework could be applied to investigate therapeutic strategies targeting immune checkpoint pathways such as CD47-SIRP*α*. Ultimately, by mathematically anchoring intracellular signaling networks to the physical mechanics of the cell membrane, this approach moves us closer to a truly predictive, multiscale understanding of immune regulation.

## Acknowledgments

Simulations were carried out at the Center for Computation and Visualization (CCV) at Brown University.

## Author contributions

Z.C. and G.E.K. conceived and designed the research. Z.C. developed and performed the computational simulations and modeling. Z.C. and N.A.D. contributed to the development of the computational framework. Z.C., N.A.D., and G.E.K. analyzed the data and interpreted the results. G.E.K. supervised the project. All authors contributed to revising the manuscript and approved the final version.

## Tables

### Dissipative particle dynamics (DPD) method and RBC model

#### Dissipative particle dynamics (DPD) method

In this work, we build our coarse-grained macrophage model and investigate the active phagocytosis process with the dissipative particle dynamics (DPD) method, a mesoscopic particle-based method that have been widely used for problems in complex fluids and soft mater (35; 23; 30; 41). In a DPD simulation, a group of atoms is represented by a bead at its center, which greatly reduces the computational cost. The interactions between two coarse-grained beads *i* and *j* are integrated into three types of forces:

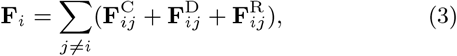

where 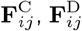 and 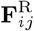 are conservative, dissipative and random forces, respectively. The typical forms of these three pairwise forces are described as follows:

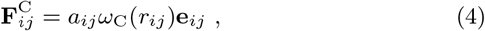

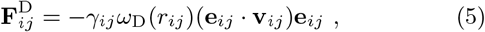

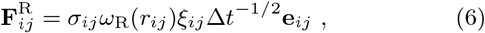

where *r*_*ij*_ is the distance between beads *i* and *j*, **e**_*ij*_ is the corresponding unit vector, and *ξ*_*ij*_ is Gaussian white noise with a mean of zero and unit variance. *a*_*ij*_ determines the strength of the conservative force, which we set to 6.0 in our simulations. *γ*_*ij*_ and *σ*_*ij*_ reflect the strength of the dissipative and random forces, respectively, and *ω*_C_(*r*_*ij*_), *ω*_D_(*r*_*ij*_), and *ω*_R_(*r*_*ij*_) are the corresponding weighting functions. To maintain the energy conservation of the simulation system and satisfy the fluctuation-dissipation theorem, two further conditions must be enforced:

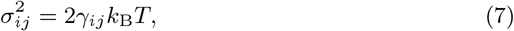

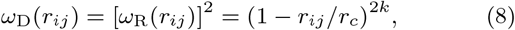

where *k*_B_ is the Boltzmann constant and *T* is the temperature. Additionally, *r*_*c*_ is the cutoff radius and *k* is the exponent for the DPD envelope.

#### RBC model

To investigate the role of each biophysical factor on the blood viscosity and, ultimately, Gaucher Disease RBC computationally, we need to model the individual RBCs accurately, without compromising the essential hydrodynamics of the system and the convective transport processes that govern the blood flow. An RBC is a highly deformable biconcave nucleus-free membrane with a diameter of ~8 *µ*m. Known as a viscoelastic object, RBC shows both liquid-like (viscous) and solid-like (elastic) responses to an applied deformation at the same time. This allows the RBCs to be very deformable to squeeze through capillaries as small as 3 *µ*m, preserve their shapes at small deformation rates, and orient in the flow direction at larger deformation rates in wider arteries. The membrane of RBC is modeled as a set of *N*_*ν*_ DPD particles with the three-dimensional coordinates *X*_*i*_ (*i* ∈ 1, …, *N*_*ν*_) in a triangulated network of springs with a dashpot on the surface. To model the incompressibility of the RBC membrane, area and volume constraints are applied. Also, considering a bending resistance between all neighboring triangles, the bending rigidity of the membrane can be mimicked.

In this work, we considered the aggregation of normal and GD RBCs under different disaggregation threshold and stiffnes, with the shear modulus and disaggregation threshold of RBCs under these conditions are summarized in Table 1. The shear modulus and bending modulus of normal RBCs in our work are selected to be *Es*_0_ = 4.792*µN/m, Eb*_0_ = 2.9 × 10^−19^*J* ((25)). We further validate our comprehensive RBC model by comparing it with experimental data, ensuring accurate simulations that capture the changes in biomechanical, rheological, and dynamic behavior of GD RBCs in response to morphological alterations and membrane stiffening (see Figure Ci to Cviii). For detailed formulas and parameters to maintain RBC morphology and mechanical properties, please refer to the Supporting Materials.

#### DPD Model Framework

In our simulations, the macrophage membrane is represented as a deformable layer composed of DPD beads, with embedded receptor slots corresponding to SIRP*α* molecules. IgG antibodies are modeled as flexible, coarse-grained beads that can bind membrane receptors. Antibody–receptor interactions are implemented through attractive interparticle potentials or reactive binding rules between IgG beads and SIRP*α* receptors. Antibody diffusion is explicitly simulated both in the 3D extracellular solution and laterally along the membrane surface after binding. These simulation outputs are used to calculate the effective SIRP*α* activation state, *X*_3_, because CD47–SIRP*α* binding is the upstream trigger of the inhibitory signaling cascade, the DPD-derived receptor occupancy provides a mechanistic proxy for the activation level of the signaling complex represented by X3. In the signaling model, *X*_3_ represents the activated CD47-SIRP*α* receptor complex. In the presence of anti-SIRP*α* antibodies, receptor occupancy reduces the availability of SIRP*α* for CD47 binding, thereby decreasing the effective activation level of *X*_3_. For anti-CD47 antibodies, *X*_3_ is inferred based on the remaining CD47 availability for SIRP*α* binding—consistent with observed increases in phagocytic activity when the SIRP*α*–CD47 interaction is blocked (83).

**Table S1.** Ordinary differential equations describing the intracellular signaling network governing RBC phagocytosis.

| State | Governing equation |
| --- | --- |
| $X_1$ | $\frac{dX_1}{dt} = -(K_7 X_1 X_{15} - K_{21} X_2) - (K_3 K_6 X_1 X_{14} - K_3 K_{20} X_{12})$ |
| $X_2$ | $\frac{dX_2}{dt} = (K_7 X_1 X_{15} - K_{21} X_2) - K_8 X_2 \left(1 - \frac{X_5}{X_5 + K_{23}}\right)^{K_{22}} + K_9 X_3$ |
| $X_3$ | $\frac{dX_3}{dt} = K_8 X_2 \left(1 - \frac{X_5}{X_5 + K_{23}}\right)^{K_{22}} - K_9 X_3$ |
| $X_4$ | $\frac{dX_4}{dt} = -K_{16} X_4 X_3 + K_{17} X_5$ |
| $X_5$ | $\frac{dX_5}{dt} = K_{16} X_4 X_3 - K_{17} X_5$ |
| $X_6$ | $\frac{dX_6}{dt} = K_{13} X_5 X_7 - K_{12} X_6 (X_{13} + X_{11} + X_{10})$ |
| $X_7$ | $\frac{dX_7}{dt} = -K_{13} X_5 X_7 + K_{12} X_6 (X_{13} + X_{11} + X_{10})$ |
| $X_8$ | $\frac{dX_8}{dt} = -K_1 K_2 K_{10} X_8 + K_{11} X_{10}$ |
| $X_9$ | $\frac{dX_9}{dt} = K_{15} X_{11} - K_4 K_5 K_{14} X_9$ |
| $X_{10}$ | $\frac{dX_{10}}{dt} = K_1 K_2 K_{10} X_8 - K_{11} X_{10}$ |
| $X_{11}$ | $\frac{dX_{11}}{dt} = -K_{15} X_{11} + K_4 K_5 K_{14} X_9$ |
| $X_{12}$ | $\frac{dX_{12}}{dt} = K_3 K_6 X_1 X_{14} - K_3 K_{20} X_{12} - K_{18} X_{12} + K_{19} X_{13}$ |
| $X_{13}$ | $\frac{dX_{13}}{dt} = K_{18} X_{12} - K_{19} X_{13}$ |
| $X_{14}$ | $\frac{dX_{14}}{dt} = -(K_3 K_6 X_1 X_{14} - K_3 K_{20} X_{12})$ |
| $X_{15}$ | $\frac{dX_{15}}{dt} = -(K_7 X_1 X_{15} - K_{21} X_2)$ |

**Table S2.** Parameters of the signaling model, their nominal values, search ranges (when applicable), and biological descriptions.

| Parameter | Value | Search range | Description |
| --- | --- | --- | --- |
| $K_1$ | $1.0 \times 10^6$ | Known | Band3 expression |
| $K_2$ | 1 | Known | Antibody |
| $K_3$ | 1 | Known | Matricellular protein TSP1 |
| $K_4$ | 1 | Known | Phosphatidyl serine (PS) |
| $K_5$ | 1 | Known | Erythrocyte rigidity |
| $K_6$ | $2.3 \times 10^{-4}$ | Known | Binding rate of CD47<sub>alt</sub> to SIRPα |
| $K_7$ | $2.3 \times 10^{-4}$ | Known | Binding rate of CD47 to SIRPα |
| $K_8$ | 0.4 | 0.3-0.5 | Activation rate of CD47-bound SIRPα |
| $K_9$ | 0.02 | — | Deactivation rate of CD47-bound SIRPα |
| $K_{10}$ | $2.0 \times 10^{-6}$ | Known | Activation rate of Fc receptor |
| $K_{11}$ | $5.0 \times 10^{-4}$ | $4.0 \times 10^{-4}$–$6.0 \times 10^{-4}$ | Deactivation rate of Fc receptor |
| $K_{12}$ | $5.2 \times 10^{-7}$ | Known | Activation rate of Myosin IIa |
| $K_{13}$ | $3.8 \times 10^{-8}$ | Known | Deactivation rate of Myosin IIa |
| $K_{14}$ | 0.51 | — | Activation rate of PS receptor |
| $K_{15}$ | $1.0 \times 10^{-3}$ | — | Deactivation rate of PS receptor |
| $K_{16}$ | 0.007 | — | Activation rate of SHP1 |
| $K_{17}$ | 0.002 | — | Deactivation rate of SHP1 |
| $K_{18}$ | 0.001 | Known | Activation rate of TSP1-bound SIRPα |
| $K_{19}$ | 0.03 | Known | Deactivation rate of TSP1-bound SIRPα |
| $K_{20}$ | 0.4 | Known | Dissociation rate of CD47<sub>alt</sub> from SIRPα |
| $K_{21}$ | 0.4 | Known | Dissociation rate of CD47 from SIRPα |
| $K_{22}$ | 3 | Known | Hill coefficient of SHP–SIRPα activation |
| $K_{23}$ | $6.0 \times 10^4$ | $5.8 \times 10^4$–$6.2 \times 10^4$ | $K_d$ coefficient of SHP–SIRPα activation |

#### Signaling Interaction Model

To characterize antibody-mediated modulation of macrophage signaling, we modeled the binding interactions between antibodies and membrane receptors such as SIRP*α* using DPD (24). In this framework, antibodies were represented as diffusing particles in the extracellular domain, capable of binding to discrete receptor sites distributed on the macrophage membrane. Binding was governed by a short-range attractive Morse potential between antibodies and receptor vertices:

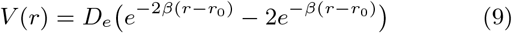

where *r* is the distance between an antibody and a receptor particle, *D*_*e*_ is the well depth (binding strength), *β* controls the interaction range, and *r*_0_ is the equilibrium binding distance. Once bound, the lateral mobility of antibodies was restricted by additional harmonic potential to mimic membrane anchoring, allowing us to capture post-binding diffusion and clustering.

**Table S3.**
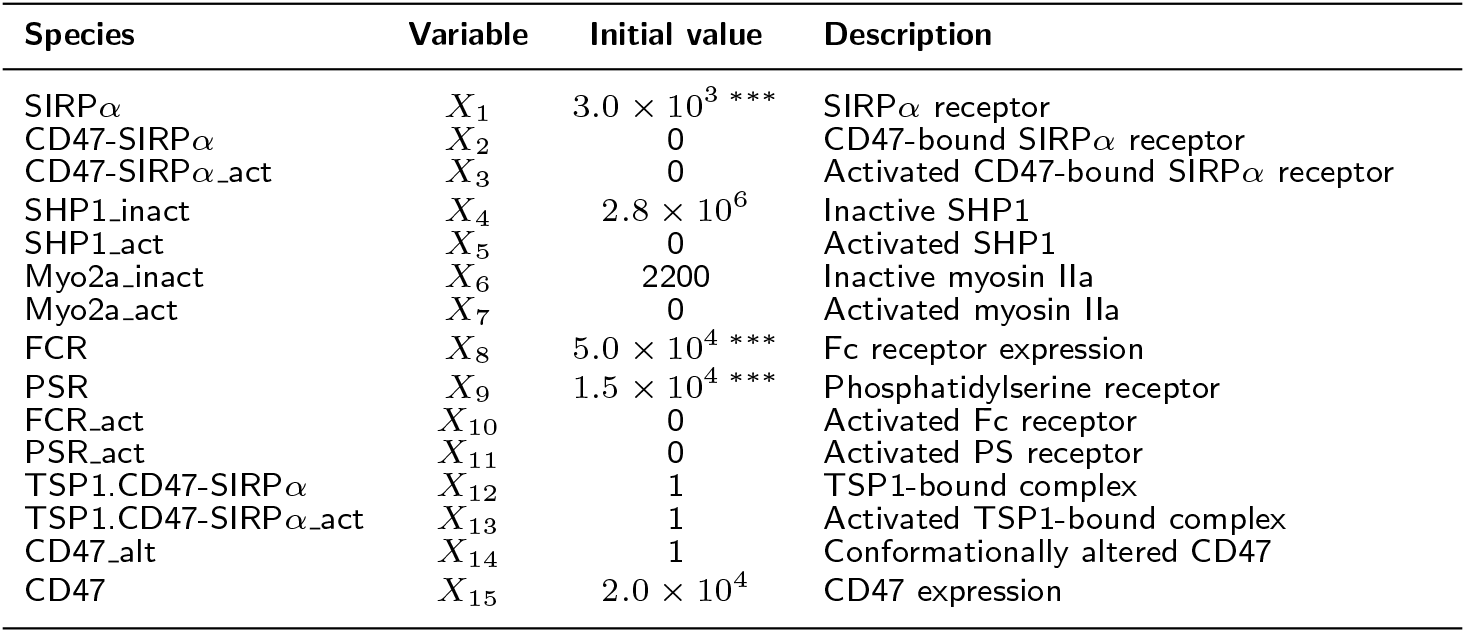
Initial conditions of the signaling model. ^∗^ Parameter obtained from fitting; ^∗∗^ Phenomenological parameters; ^∗∗∗^ Macrophage receptor levels.

**Table S4.** Comparison of simulated and experimental diffusion coefficients. Physical values are obtained using the DPD unit mapping from Table S1.

| Sample | $D_{\text{DPD}}$ | $D_{\text{phys}}$ (cm<sup>2</sup>/s) |
| --- | --- | --- |
| Green Line | $1.46 \times 10^{-3}$ | $1.05 \times 10^{-7}$ |
| Blue Line | $5.07 \times 10^{-4}$ | $3.66 \times 10^{-8}$ |
| Black Line | $1.83 \times 10^{-4}$ | $1.32 \times 10^{-8}$ |
| IgG (experimental) (27; 58; 81) | — | $10^{-8}$–$10^{-6}$ |

The spatial layout of receptor sites was initialized with non-overlapping positions across the membrane surface. Receptor occupancy over time was recorded as the fraction of bound receptors, denoted *X*_3_(*t*), and was extracted directly from the DPD simulations under different molecular crowding and antibody concentration conditions. The resulting *X*_3_(*t*) profiles were interpolated and used as input or constraints in the PINNs-based ODE model of CD47-SIRP*α* signaling. This hybrid DPD-ODE approach links antibody binding dynamics to macrophage activation states and enables evaluation of immunomodulatory strategies targeting the CD47-SIRP*α* axis (60).

### DPD Simulation Parameters for Antibody–Receptor Binding

#### Particle Types and Initial Configuration

Due to the limitations of experimental *in vitro* measurements mentioned earlier, we employed computational approaches. Advances in computational power over the past two decades have stimulated the development and application of multiscale biophysical models for studying complex cellular systems (63; 70; 10; 13; 12). In parallel, machine learning approaches have increasingly been applied to collective cell dynamics and particle-based simulations (88; 87). A coarse-grained Dissipative Particle Dynamics (DPD) model was employed to simulate antibody–receptor interactions occurring on the macrophage membrane. The simulated system consisted of three main particle types representing the extracellular environment, antibodies, and membrane structures. Solvent particles were used to represent the surrounding extracellular fluid and served as a thermalized background that mediates hydrodynamic interactions between all components of the system. Antibodies, such as anti-SIRP*α* IgG molecules, were modeled as spherical DPD beads with an effective diameter of *σ* = 1.0 *µm*. These particles were initially distributed randomly throughout the solvent volume and allowed to diffuse freely until encountering receptor sites on the macrophage surface.

The macrophage membrane was represented as a triangulated network of DPD particles forming a deformable surface. Receptor sites corresponding to SIRP*α* molecules were implemented as a subset of these membrane particles. These receptors were arranged in predefined spatial clusters to mimic the presence of lipid raft–like membrane domains that locally enrich receptor density. In the initial configuration, 300 antibody particles were uniformly distributed within the bulk solvent domain. The macrophage membrane consisted of approximately 1000 DPD beads forming a spherical cap geometry, among which 200 particles were designated as receptor sites. Receptor clusters occupied roughly 30% of the membrane area in order to reproduce experimentally observed membrane microdomain organization.

#### Interaction Potentials

Binding interactions between antibody particles and SIRP*α* receptors were modeled using a Morse potential. To prevent unphysical particle overlap and to maintain the hydrodynamic consistency of the system, standard DPD conservative, dissipative, and random forces were applied between all particle pairs. The conservative force followed the standard soft repulsive form

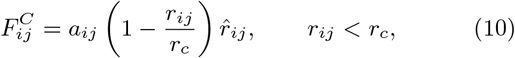

with a maximum repulsion parameter *a*_*ij*_ = 25 and a cutoff radius *r*_*c*_ = 1.0 *σ*. After binding occurs, antibody particles were additionally connected to receptor sites through a harmonic spring potential to represent the reduced lateral mobility of membrane-bound complexes.

#### Simulation Domain and Conditions

The simulation domain had dimensions of 60 × 60 × 50 *µm*^3^. Periodic boundary conditions were imposed along the *x* and *z* directions, while reflective boundaries were applied along the *y* direction to mimic confinement near the membrane interface. The simulations were typically run for 4 × 10^6^ to 8 × 10^6^ time steps using a modified velocity–Verlet integration scheme combined with a DPD thermostat to maintain the system temperature.

**Table S5.**
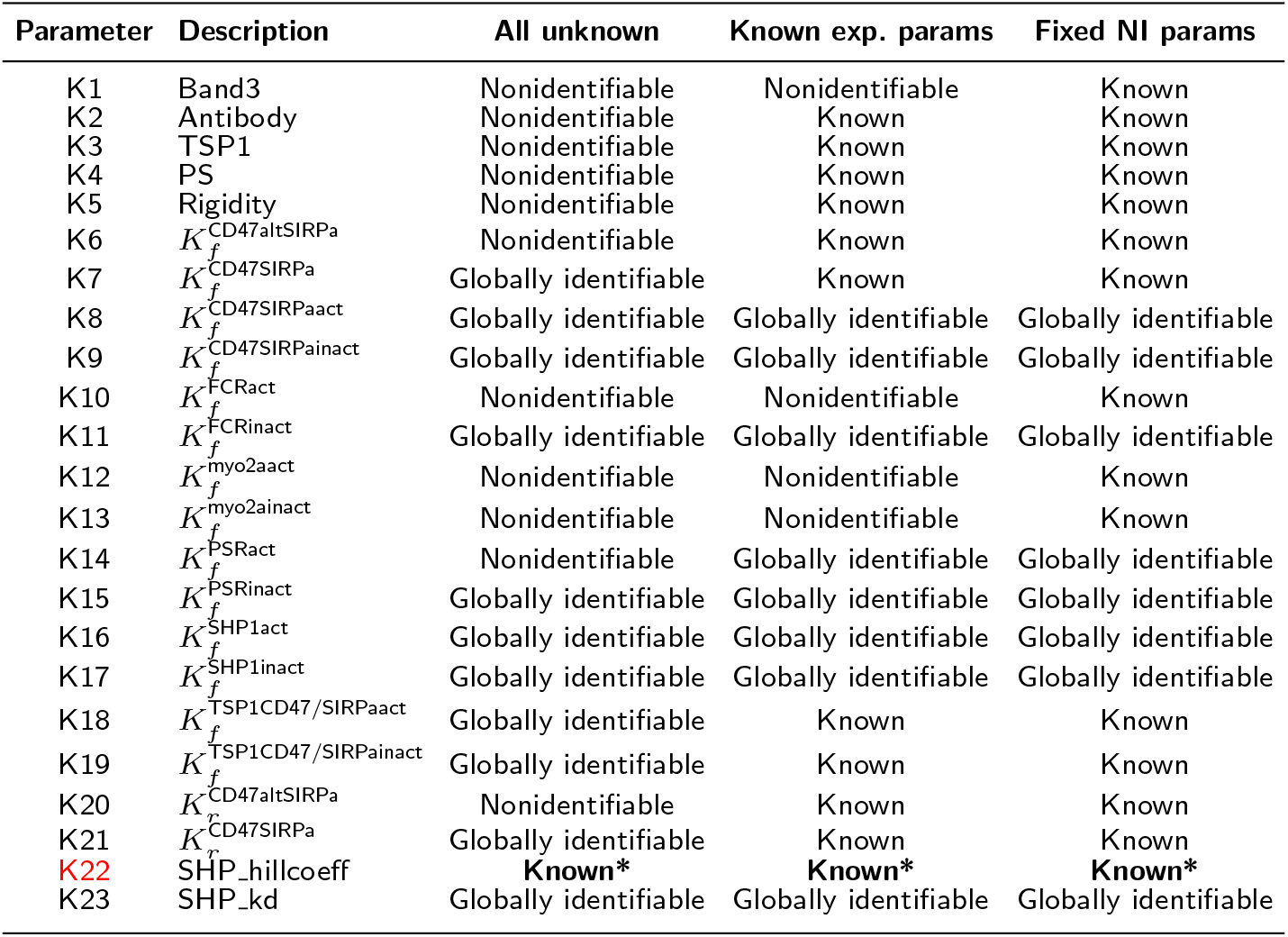
Summary of structural identifiability results for the signaling model parameters under three scenarios: (i) all parameters treated as unknown, (ii) selected parameters fixed based on available experimental data, and (iii) additional nonidentifiable parameters fixed according to prior biological knowledge. “**Known\***” indicates parameter *K*_22_, which was fixed to its experimentally determined value (see Materials and Methods for details).

**Table S6.**
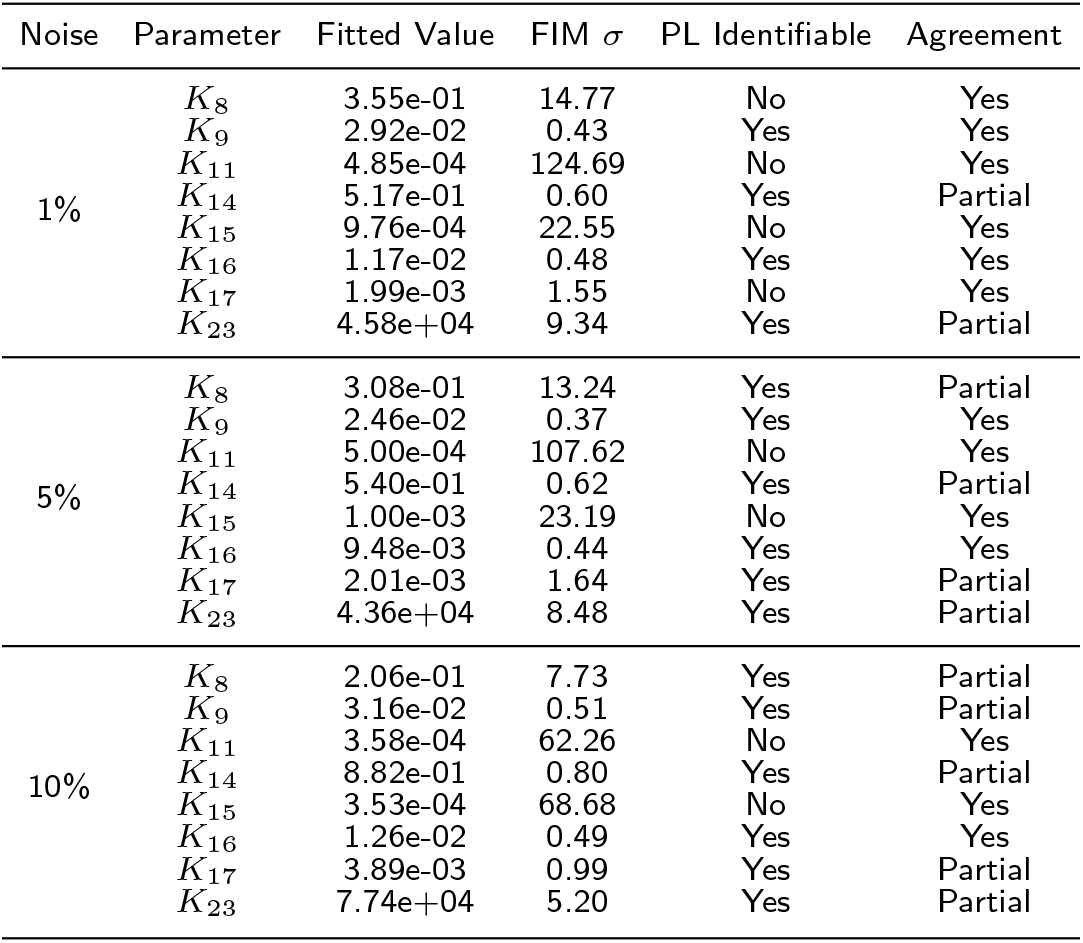
Summary of practical identifiability results across observational noise levels. FIM *σ* denotes the relative standard deviation estimated from the inverse Fisher Information Matrix. PL indicates whether the profile likelihood exceeded the 95% confidence threshold. Agreement reflects consistency between FIM and profile likelihood conclusions.

| Noise | Parameter | Fitted Value | FIM $\sigma$ | PL Identifiable | Agreement |
| --- | --- | --- | --- | --- | --- |
| 1% | $K_8$ | 3.55e-01 | 14.77 | No | Yes |
| | $K_9$ | 2.92e-02 | 0.43 | Yes | Yes |
| | $K_{11}$ | 4.85e-04 | 124.69 | No | Yes |
| | $K_{14}$ | 5.17e-01 | 0.60 | Yes | Partial |
| | $K_{15}$ | 9.76e-04 | 22.55 | No | Yes |
| | $K_{16}$ | 1.17e-02 | 0.48 | Yes | Yes |
| | $K_{17}$ | 1.99e-03 | 1.55 | No | Yes |
| | $K_{23}$ | 4.58e+04 | 9.34 | Yes | Partial |
| 5% | $K_8$ | 3.08e-01 | 13.24 | Yes | Partial |
| | $K_9$ | 2.46e-02 | 0.37 | Yes | Yes |
| | $K_{11}$ | 5.00e-04 | 107.62 | No | Yes |
| | $K_{14}$ | 5.40e-01 | 0.62 | Yes | Partial |
| | $K_{15}$ | 1.00e-03 | 23.19 | No | Yes |
| | $K_{16}$ | 9.48e-03 | 0.44 | Yes | Yes |
| | $K_{17}$ | 2.01e-03 | 1.64 | Yes | Partial |
| | $K_{23}$ | 4.36e+04 | 8.48 | Yes | Partial |
| 10% | $K_8$ | 2.06e-01 | 7.73 | Yes | Partial |
| | $K_9$ | 3.16e-02 | 0.51 | Yes | Partial |
| | $K_{11}$ | 3.58e-04 | 62.26 | No | Yes |
| | $K_{14}$ | 8.82e-01 | 0.80 | Yes | Partial |
| | $K_{15}$ | 3.53e-04 | 68.68 | No | Yes |
| | $K_{16}$ | 1.26e-02 | 0.49 | Yes | Yes |
| | $K_{17}$ | 3.89e-03 | 0.99 | Yes | Partial |
| | $K_{23}$ | 7.74e+04 | 5.20 | Yes | Partial |

#### Estimation and Validation of Diffusion Coefficients

In the diffusive regime, the MSD grows linearly with time according to the Einstein relation for three–dimensional Brownian motion,

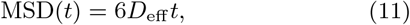

where *D*_eff_ denotes the effective diffusion coefficient. Linear regression was therefore applied to the long-time portion of the MSD curves, and the slope divided by six to obtain the diffusion coefficient in DPD units, *D*_DPD_. In addition, Figure S1D shows the statistical distribution of diffusion coefficients estimated from individual particle trajectories. To convert the simulated diffusion coefficients to physical units, we employed the unit mapping of the DPD model (Table S1). The characteristic length and time scales are *L*^∗^ = 1.0 × 10^−6^ m and *t*^∗^ = 1.386 × 10^−4^ s, which yield the conversion factor

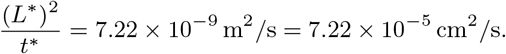

Accordingly, the physical diffusion coefficient is obtained as

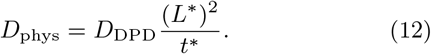

Using this mapping, the diffusion coefficients extracted from the MSD curves were converted to physical units. For example, the green trajectory in Figure S1C yields *D*_DPD_ = 1.46 × 10^−3^, corresponding to *D*_phys_ = 1.05 × 10^−7^ cm^2^*/*s. The full set of estimated diffusion coefficients is summarized in Table S4.

The simulated values fall within the experimentally reported diffusion range for IgG antibodies (10^−8^–10^−6^ cm^2^*/*s) (27; 58; 81). This validation is particularly relevant for modeling antibody-mediated processes in diseases such as GD, which is characterized by elevated levels of circulating IgG autoantibodies (including IgG1, IgG2, and IgG3) (56). In particular, the higher diffusivity observed for the green and blue trajectories reflects the smaller effective particle sizes represented in the coarse-grained DPD model, whereas the black trajectory provides a lower bound consistent with restricted transport in crowded environments. The overall agreement with experimental measurements supports the validity of the DPD framework for capturing mesoscale antibody diffusion.

**Table S7.** Comparison of PINNs parameter estimation with and without search space constraints. Search limits were applied to parameters identified as weakly constrained in the practical identifiability analysis (red). Constraining the search space reduces variance and improves stability without affecting well-identified parameters.

| Parameter | True Value | Without Limit |  |  | With Limit |  |  |
| --- | --- | --- | --- | --- | --- | --- | --- |
|  |  | Mean | Std | Median | Mean | Std | Median |
| K8 | 4.00e-1 | 7.16e-1 | 1.42e-1 | 7.74e-1 | 3.83e-1 | 3.37e-2 | 3.7e-1 |
| K9 | 2.00e-2 | 1.74e-2 | 4.39e-4 | 1.75e-2 | 1.98e-2 | 3.49e-4 | 1.97e-2 |
| K11 | 5.00e-4 | 1.24e-3 | 7.02e-4 | 8.90e-4 | 5.15e-4 | 9.00e-6 | 5.10e-4 |
| K14 | 5.10e-1 | 5.09e-1 | 3.71e-3 | 5.10e-1 | 5.21e-1 | 2.71e-4 | 5.21e-1 |
| K15 | 1.00e-3 | 9.12e-4 | 2.43e-4 | 1.00e-3 | 1.11e-3 | 3.3e-5 | 1.10e-3 |
| K16 | 7.00e-3 | 6.92e-3 | 1.42e-4 | 6.90e-3 | 6.88e-3 | 3.60e-5 | 6.87e-3 |
| K17 | 2.00e-3 | 1.95e-3 | 1.08e-4 | 1.97e-3 | 1.94e-3 | 2.20e-5 | 1.93e-3 |
| K23 | 6.00e+4 | 4.10e+4 | 5.82e+3 | 4.23e+4 | 5.92e+4 | 1.41e+3 | 5.85e+4 |

**Table S8.** Relative error (%) of inferred parameters under different noise levels for PINNs and tanh-cPIKANs.

| Noise | PINNs |  |  |  |  |  |  |  |  | Tanh-cPIKANs |  |  |  |  |  |  |  |
| --- | --- | --- | --- | --- | --- | --- | --- | --- | --- | --- | --- | --- | --- | --- | --- | --- | --- |
|  | K8 | K9 | K11 | K14 | K15 | K16 | K17 | K23 |  | K8 | K9 | K11 | K14 | K15 | K16 | K17 | K23 |
| 0% | 7.82 | 1.86 | 0.76 | 0.05 | 0.18 | 1.99 | 2.65 | 2.43 |  | 7.16 | 2.42 | 12.50 | 0.63 | 5.17 | 1.44 | 0.73 | 1.92 |
| 5% | 7.96 | 2.06 | 2.43 | 2.22 | 1.01 | 1.78 | 2.98 | 2.27 |  | 8.80 | 2.61 | 13.11 | 1.25 | 5.78 | 1.73 | 1.10 | 2.30 |
| 10% | 9.90 | 7.00 | 0.89 | 1.95 | 0.54 | 2.71 | 5.64 | 1.15 |  | 9.50 | 3.09 | 12.82 | 1.50 | 5.94 | 2.19 | 1.47 | 2.73 |

To interpret the coarse-grained representation in molecular terms, we estimated the number of physical molecules represented by each DPD particle using the concentration mapping defined in Table S1. The concentration unit was chosen as *C*^∗^ = 1.0 nM, corresponding to a number density *n*^∗^ = *C*^∗^*V* ^∗^*/ρ*, where *V* ^∗^ = (*L*^∗^)^3^ is the volume unit and *ρ* is the particle number density. For an antibody-like species initialized at *C*_0_ = 1 nM (10^−3^ mol*/*m^3^), the number of moles represented by each DPD particle is approximately

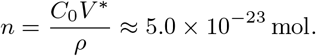

Multiplying by Avogadro’s number yields approximately 30 molecules per DPD particle. Thus, each coarse-grained particle in the simulation effectively represents 𝒪(10^1^) antibody molecules, providing a physically interpretable mapping between the DPD representation and the underlying molecular population.

#### Validation of Antibody Dynamics

As shown in Figure S2A, our simulation predicts a time-resolved receptor occupancy curve where the activation of the CD47–SIRP*α* complex (*X*_3_) reaches a plateau of approximately 70% of the healthy baseline (90). This simulated 30% reduction is consistent with experimental observations indicating a diminished “don’t eat me” signal in RBCs, providing a quantitative basis to evaluate how CD47 depletion or receptor blockade impacts inhibitory signaling activity (39).

The binding kinetics extracted from the early-time regime correspond to an effective on-rate of 1.2 × 10^5^ M^−1^s^−1^, comparable to surface plasmon resonance measurements of CD47–SIRP*α* binding, which fall in the range of 10^5^–10^6^ M^−1^s^−1^ (75). Figure S2B depicts the spatial distribution of antibody–receptor complexes across the macrophage membrane. The simulations establish a quantitative framework for analyzing diffusion and localized binding dynamics. The results are consistent with fluorescence microscopy observations, where receptor–ligand complexes organize into microdomains with elevated local densities (74). While these results support the biological relevance of the model, it is important to note that the current framework does not include an explicit cytoskeletal scaffold. Such structures are known to restrict receptor diffusion and organize membrane domains in red blood cells and neurons (44; 45; 89; 10; 13), and their absence may contribute to quantitative differences between simulation and experiment.

#### Structural and Practical Identifiability Ana

When all parameters were treated as unknown, several kinetic rates were found to be globally identifiable, including *K*_7_, *K*_8_, *K*_9_, *K*_11_, *K*_15_, *K*_16_, *K*_17_, *K*_18_, *K*_19_, *K*_21_, and *K*_23_. These parameters are structurally recoverable from perfect measurements and therefore form the core set of candidates for subsequent inverse analysis. In contrast, parameters associated with membrane composition, rigidity, alternative receptor binding pathways, and certain cytoskeletal activation steps, such as *K*_1_–*K*_6_, *K*_10_, *K*_12_, *K*_13_, and *K*_20_, were determined to be structurally nonidentifiable. Their nonidentifiability arises from model symmetries and compensatory transformations that leave the observable outputs invariant.

Incorporating experimentally known parameters into the analysis significantly improved identifiability of specific reaction rates. In particular, fixing biologically well-characterized quantities (e.g., ligand concentrations or measured kinetic constants) resolved several previously nonidentifiable directions in parameter space. Under this partially constrained setting, additional parameters, including *K*_14_, became globally identifiable. This demonstrates that identifiability in nonlinear biochemical networks depends strongly on prior knowledge and observable selection.

In the final reduced scenario, parameters classified as structurally nonidentifiable and lacking sufficient experimental constraints were fixed to representative literature values. This identifiability-informed reduction eliminates symmetry-induced ambiguities and removes flat directions from the parameter space. The resulting reduced parameter set—comprising *K*_8_, *K*_9_, *K*_11_, *K*_14_, *K*_15_, *K*_16_, *K*_17_, and *K*_23_—forms the structurally identifiable subset used in subsequent inverse modeling and PIN-based parameter estimation.

Notably, the condition number decreased slightly at the highest noise level (10%), reflecting changes in local curvature near the perturbed optimum rather than improved identifiability. Strong positive coupling is consistently observed between *K*_8_ and *K*_23_, as well as between *K*_9_ and *K*_16_, while *K*_8_ and *K*_16_ exhibit pronounced negative correlation. These dominant dependencies persist under increasing noise, indicating intrinsic parameter coupling in the model structure rather than noise-induced artifacts. In contrast, parameters such as *K*_11_ and *K*_15_ do not display strong pairwise correlations but instead exhibit weak curvature signatures in the local sensitivity landscape. This weak curvature indicates that the objective function varies only marginally with respect to these parameters near the optimum, which results in broad, shallow minima and large uncertainty estimates. Such flat directions in the loss surface are consistent with their flat profile likelihoods and confirm their practical non-identifiability despite structural identifiability.

Profile likelihood analysis provides complementary global insight. At 1% noise, parameters *K*_9_, *K*_14_, *K*_16_, and *K*_23_ exhibit finite confidence intervals exceeding the 95% *χ*^2^ threshold, indicating practical identifiability. In contrast, *K*_11_, *K*_15_, and *K*_17_ fail to cross the threshold, demonstrating flat likelihood profiles and non-identifiability. At 5% noise, a similar pattern is observed, with *K*_8_ becoming identifiable while *K*_11_ and *K*_15_ remain unidentifiable. At 10% noise, *K*_8_, *K*_9_, *K*_14_, *K*_16_, *K*_17_, and *K*_23_ retain identifiable profiles, whereas *K*_11_ and *K*_15_ persistently exhibit flat profiles with effectively unbounded confidence intervals.

Parameters *K*_16_ and *K*_9_ consistently demonstrate strong curvature and bounded likelihood profiles across noise levels that indicate robust recoverability. Conversely, *K*_11_ and *K*_15_ exhibit large FIM-derived variance estimates and flat profile likelihoods, confirming practical non-identifiability despite being structurally identifiable. For practical identifiability assessment, two complementary criteria were used. The FIM was computed in log-parameter space, and uncertainty was quantified as 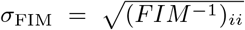. Parameters with *σ*_FIM_ *<* 0.5 were classified as locally identifiable, while parameters with 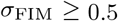 were considered weakly constrained. Because *σ*_FIM_ is defined in log space, this threshold corresponds approximately to less than 65% multiplicative uncertainty in parameter magnitude. Profile likelihood identifiability was evaluated using the 95% *χ*^2^ threshold for one degree of freedom. Since the objective function was defined as 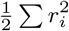, the corresponding cutoff for practical identifiability was 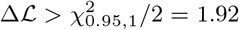. Agreement between FIM and profile likelihood was defined as consistency between these two classifications. We demonstrate that parameter identifiability is constrained by strong correlations predicted by the FIM and confirmed via Monte Carlo sampling S4.

#### Comparison of *q*-Scheduling Methods

An additional component of the modeling framework involved a *q*-scheduling strategy designed to improve the convergence and stability of PINNs. In this formulation, the parameter *q* determines the relative weight between the physics-based loss term and the data-based loss term in the composite training objective. Gradually increasing the contribution of the physics loss during training allows the neural network to first approximate the available experimental data before enforcing the full system of governing equations. This approach is conceptually analogous to temperature scheduling methods used in thermostats such as Berendsen or Nosé–Hoover, which regulate the rate at which systems approach equilibrium (78). By preventing stiff ODE residuals from dominating the early stages of training, the scheduling strategy reduces the risk of convergence to poor local minima and improves the stability of parameter inference. Comparative tests across different scheduling approaches indicated that smooth and continuous *q*-adjustment generally yields better training stability and prediction accuracy than stepwise schemes (84).

In addition to loss weighting, we explored how the number and distribution of residual points affect the performance of PINNs in inverse parameter estimation. We compared both fixed and randomized residual sampling strategies across a range of residual point counts. Our results show that model accuracy is highly sensitive to this configuration: using a moderate number of residual points—neither too sparse nor overly dense—leads to significantly improved parameter recovery. Furthermore, randomized sampling consistently outperformed fixed sampling at comparable point counts, likely due to better generalization and reduced bias in residual enforcement. This behavior is consistent with recent work showing that residual point refinement and adaptive sampling act as a form of curriculum learning, guiding the network toward physically consistent solutions while avoiding optimization instability (82; 52). These findings emphasize that residual point design is a key component in building robust PINNs architectures for biological inference problems.

We systematically evaluated different *q*-scheduling strategies and their impact on PINNs parameter inference accuracy. The parameter *q* controls the relative weight of the physics-based loss term during training, and scheduling its evolution over iterations has been proposed as a way to stabilize optimization and improve convergence (86; 78). Figure S1, panels A and B, show the evolution of *q* as a function of iteration for several scheduling strategies, targeting final values of *q* = 2 in panel A and *q* = 10 in panel B. Among the methods tested, the Berendsen-type schedules (8) and the Nose–Hoover-inspired schedule (53) achieve smooth and gradual increases in *q*, whereas the step method exhibits abrupt jumps. Smooth convergence of *q* is desirable because it helps avoid the premature dominance of the physics loss and allows the model to fit the sparse data during the early stages of training (38). The baseline case (*q* = 1, with no warm-up) exhibited relatively poor inference performance, and step scheduling likewise showed unfavorable behavior across the tested target *q* values. By contrast, Berendsen-based and Nose–Hoover schedules yielded qualitatively better results overall. In particular, the Berendsen-type methods displayed the most favorable performance and consistently outperformed both the linear and step strategies at *q* = 2 and *q* = 10. Collectively, these observations suggest that thermostat-inspired warm-up schedules can improve training stability and support better identifiability.

**Table S9.** DPD and fluid membrane parameters in phagocytosis simulations.

| Parameter | Symbol | Value used |
| --- | --- | --- |
| strength of the conservative force | $a_{ij}$ | 6 |
| strength of the dissipative force | $\gamma_{ij}$ | 15 |
| strength of the random force | $\sigma_{ij}$ | 2.63 |
| cutoff distance of DPD interaction | $r_c$ | 2.7 |
| exponent of DPD interaction | $k$ | 0.5 |
| temperature | $k_B T$ | 0.23 |
| energy unit of fluid membrane interaction | $\varepsilon$ | 1.0 |
| length unit of fluid membrane interaction | $\sigma$ | 1.0 |
| cutoff distance of fluid membrane interaction | $r_{cut}$ | 2.6 |
| exponent of fluid membrane interaction | $\zeta$ | 4 |
| angular coefficient of fluid membrane interaction | $\mu$ | 3 |
| spontaneous angle of fluid membrane interaction | $\sin \theta_0$ | 0 |

It is noteworthy that the step scheduling results are identical for *q* = 2 and *q* = 10. This is because, in both cases, the model selected the best checkpoint at approximately 18,000 iterations—before the scheduled step increase in *q* took effect. Consequently, the later jump in *q* inherent to the step method had no impact on the final reported results. This observation highlights a potential drawback of step scheduling: its delayed activation of the physics loss can lead the model to overfit the early data-fitting phase and ignore the later dynamics. These results highlight the importance of warm-up scheduling when employing PINNs for inverse problems with stiff physics constraints. In particular, Berendsen-type *q*-scheduling provides a practical and effective approach to mitigate training pathologies and enhance inference accuracy (38; 86).

#### Physics-Informed Neural Networks Architecture

In the baseline PINNs model, the network is a fully connected feed-forward neural network with tanh activation functions. For an input *t*, the forward propagation through an *L*-layer network is given by

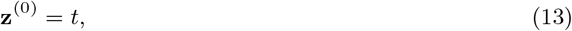

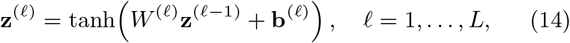

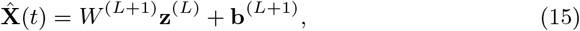

where *W* ^(*ℓ*)^ and **b**^(*ℓ*)^ denote the weight matrices and bias vectors of layer *ℓ*. The final layer is linear to allow unrestricted output scaling.

All unknown kinetic parameters are optimized jointly with the network weights.

#### Physics-Informed Kolmogorov–Arnold Networks Architecture

To investigate whether structured representations improve inverse robustness, we replace the MLP with a Kolmogorov– Arnold Network (KAN) (47). KANs are motivated by the Kolmogorov–Arnold representation principle, which approximates multivariate functions via compositions of univariate nonlinear mappings rather than affine transformations.

In the Chebyshev-based variant (cPIKAN) (66; 68), the learnable univariate mappings are parameterized using Chebyshev polynomials:

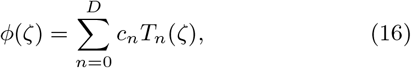

where *T*_*n*_(*ζ*) denotes the *n*-th Chebyshev polynomial and {*c*_*n*_} are trainable coefficients.

In this work, we adopt a modified architecture referred to as *tanh-cPIKAN*, which introduces nested tanh nonlinearities to improve numerical stability. Let *σ*(*x*) = tanh(*x*). For an *L*-layer network, the forward mapping is defined recursively as

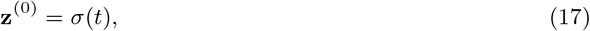

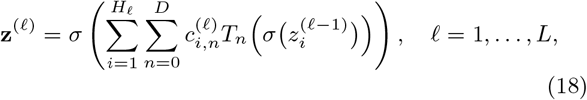

with final linear readout

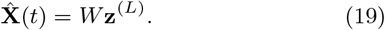

The linear readout allows output scaling via *W* while preserving bounded intermediate representations throughout the network. Beyond bounding activations, the tanh nonlinearity modifies the optimization geometry. Let **z**(*θ*) denote the Chebyshev expansion and define the contracted representation

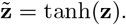

By the chain rule, the parameter Jacobian becomes

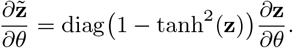

**Figure S1.**
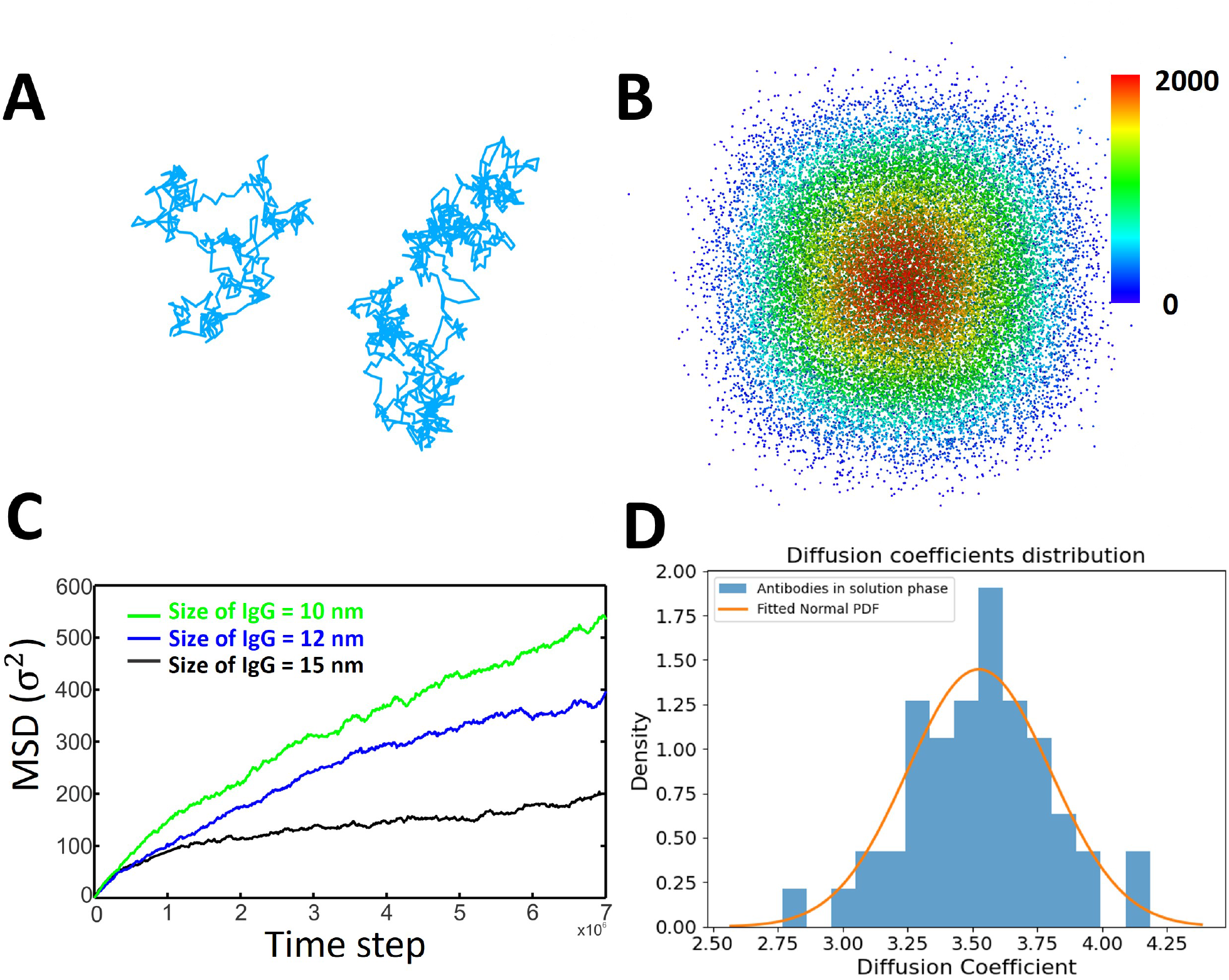
Validation of diffusive behavior in DPD simulations. (A) Sample trajectories of selected particles undergoing diffusion in 3D space, demonstrating characteristic random-walk behavior. (B) Radial particle density distribution centered at the origin, with color mapping from blue (low) to red (high) density, showing isotropic and spherically symmetric spreading consistent with Brownian diffusion. (C) Mean squared displacement (MSD) of IgG of different sizes (10, 12, and 15 nm), where smaller IgG exhibit faster diffusion as indicated by steeper MSD curves. (D) Distribution of diffusion coefficients for IgG in the solution phase (blue histogram), fitted with a normal probability density function (orange line), illustrating variability in antibody mobility across the ensemble.

Thus, tanh inserts a diagonal contraction matrix with entries in (0, 1] into the Jacobian.

For the squared residual loss ℒ(*θ*) = ∥*r*(*θ*)∥^2^, the dominant curvature term is the Gauss–Newton matrix *H*_*GN*_ = *J*^⊤^*J*. With tanh activations, this becomes

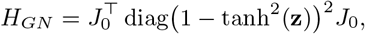

where *J*_0_ denotes the Jacobian of the polynomial expansion without contraction.

Consequently, tanh rescales curvature directions and attenuates those associated with large-magnitude activations. In Chebyshev-based networks, high-degree polynomial interactions may amplify derivatives and induce sharp curvature in the loss landscape. The repeated contraction induced by tanh mitigates this effect by limiting activation growth and reducing curvature anisotropy, thereby improving numerical stability during physics-informed training (40). For first-order optimizers such as Adam, this contraction promotes more balanced gradient magnitudes and reduces oscillatory behavior caused by high-frequency components.

Both the PINNs and PIKANs architectures are trained using the identical composite physics-informed loss; only the functional representation differs.

#### Pairwise scatter plots of Monte Carlo parameter estimates

To evaluate the variability and correlations among trainable parameters, we conducted 100 independent Monte Carlo (MC) replicates of the parameter estimation procedure and visualized the results in a scatter matrix (Figure S4). Each off-diagonal panel displays the joint distribution between two parameters, while the diagonal panels show the self-correlation of each parameter.

The scatter plots reveal distinct correlation structures across several parameter pairs. For instance, strong negative correlations are observed between K8–K17 and K8–K23, whereas K14–K15 exhibit a pronounced positive correlation. Other parameter pairs show weaker or no clear dependency, reflecting partial independence. These empirical trends align well with the Fisher Information Matrix (FIM)-based correlation predictions, thereby reinforcing the robustness of the identifiability analysis. Overall, the scatter matrix provides an intuitive visualization of parameter coupling and supports the conclusions drawn from the analytical FIM approach.

**Figure S2.**
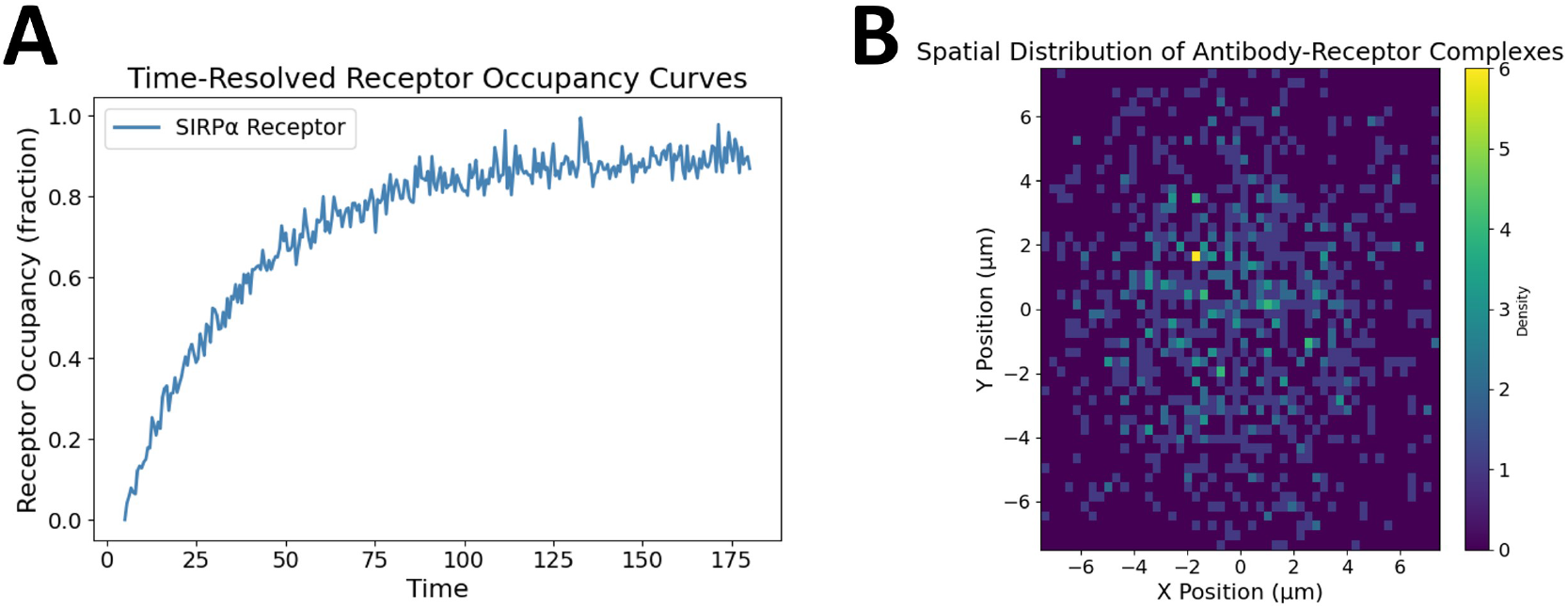
Simulation-derived observables characterizing IgG antibody dynamics and membrane engagement. (A) Time-resolved receptor occupancy curve for anti-SIRP*α* IgG, showing progressive binding to SIRP*α* receptors on the macrophage membrane. (B) Spatial distribution heatmap of antibody–receptor complexes on the membrane, illustrating clustering due to lateral diffusion and binding kinetics.

**Figure S3.**
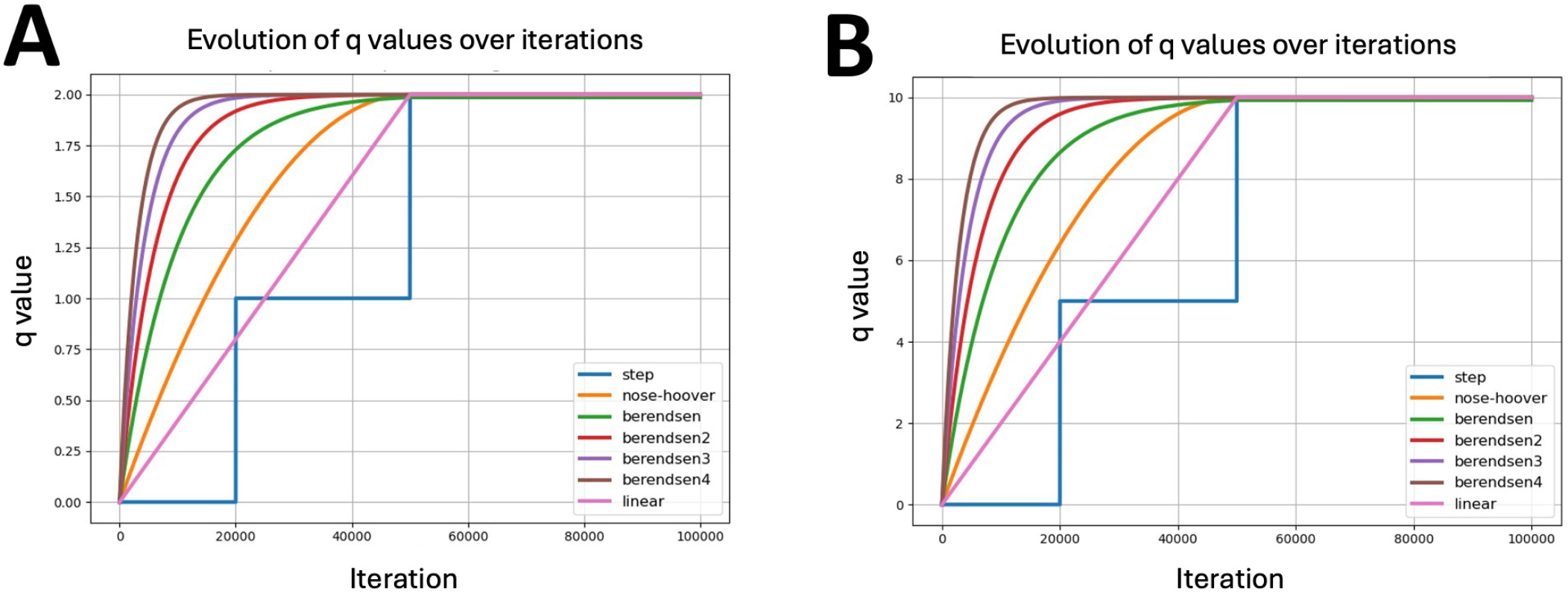
Comparison of the evolution of *q* values across *q*-scheduling methods. (A) Evolution of *q* values over iterations using various scheduling strategies with a target value of *q* = 2. (B) Same as in (A), but for *q* = 10. Berendsen-type methods show smoother convergence and outperform step scheduling.

#### Effect of Collocation Strategy: Fixed vs. Random Sampling

To further evaluate the training robustness and sensitivity to the collocation point configuration, we analyzed the performance of both PINNs and Tanh-cPIKANs using two types of physics-based residual sampling strategies: (i) **fixed collocation points** (denoted by **F**), and (ii) **randomly resampled collocation points during training** (denoted by **R**). Here, F100 refers to training with 100 fixed collocation points, while R1000 denotes resampling 1000 collocation points at each epoch. The relative parameter estimation errors for PINNs and Tanh-cPIKANs across these configurations are summarized in Tables S10 and S11, respectively.

Our findings reveal several important trends:

1. **Increasing the number of collocation points does not guarantee improved performance** In both PINNs and Tanh-cPIKANs, increasing the number of fixed collocation points beyond a certain threshold (e.g., F1000 or F1500) does not lead to consistently lower parameter inference error. For some parameters, performance plateaus or even worsens. It highlights the diminishing returns and potential over-constraining effect of excessive residual points.
2. **Random resampling can improve generalization at low point counts, but its benefits diminish with more collocation points** Comparing R100 and R1000 to their fixed counterparts (F100 and F1000), we observe that random resampling often leads to lower variance and better accuracy at lower point counts. However, as the number of collocation points increases, the advantage of stochastic sampling reduces, and in some cases the performance becomes comparable or slightly inferior to fixed strategies. This suggests that the benefits of randomization are more pronounced in lower number of collocation points.
3. **Tanh-cPIKANs exhibit greater robustness across collocation strategies, particularly under noise and limited residual supervision** Regardless of whether fixed or random sampling is used, and regardless of the number of collocation points, Tanh-cPIKANs achieve lower relative error in estimating most parameters. This robustness can be attributed to their hierarchical representation and localized basis structure, which allows for more flexible adaptation to the physical constraints even with limited or noisy residual supervision.
4. **INNs are more sensitive to noise and sampling choices** In the PINNs results, several parameters—such as *K*_8_, *K*_9_, and *K*_15_—exhibit error increases of up to fourfold when transitioning from 0% to 10% noise, and training performance varies significantly across collocation strategies. In contrast, Tanh-cPIKANs maintain stable performance under the same conditions.
5. **Random sampling does not uniformly outperform fixed sampling** Although random sampling has theoretical benefits in reducing overfitting to specific collocation locations and improving generalization, our experiments show that this is not universally the case. For both PINNs and PIKANs, certain parameters (e.g., *K*_14_, *K*_16_) show minimal sensitivity to the sampling method, while others show fluctuating gains. This indicates that the effectiveness of random sampling is model- and problem-dependent. In summary, these results highlight the nuanced trade-offs in collocation point strategies. While increasing collocation points or using random resampling can help under certain conditions, these factors alone are not sufficient to guarantee improved parameter recovery. Instead, architectural choices—such as those employed in tanh-cPIKANs—play a more critical role in ensuring robustness and accuracy in inverse problem settings.

**Figure S4.**
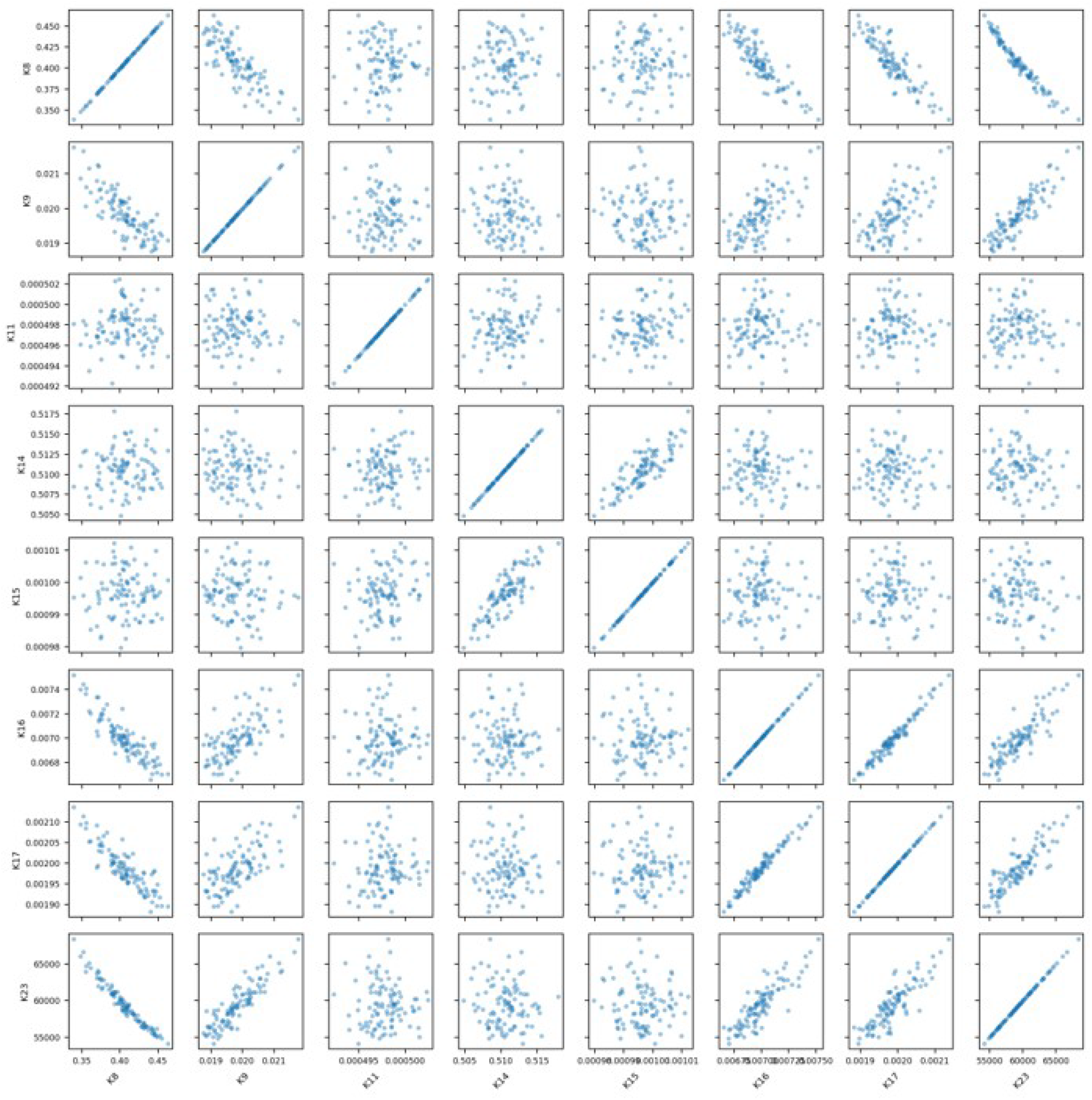
Pairwise scatter plots of Monte Carlo parameter estimates. Scatter plots of estimated parameters (K8, K9, K11, K14, K15, K16, K17, K23) from 100 Monte Carlo replicates. Each panel shows joint distributions, while diagonal panels show marginal distributions. Both positive and negative correlations are observed, consistent with FIM-based analysis.

**Table S10.** PINNs: Relative parameter estimation errors (%) under fixed (F) and randomly resampled (R) collocation point configurations. F-*N* denotes training with *N* fixed collocation points, while R-*N* denotes resampling *N* points at each epoch. Errors are computed relative to nominal parameter values after convergence. Entries in **bold** indicate the most accurate configuration (lowest relative error) for each parameter across sampling strategies and Network architecture.

| Params | F-100 | F-200 | F-1000 | F-1500 | R-100 | R-200 | R-1000 | R-1500 |
| --- | --- | --- | --- | --- | --- | --- | --- | --- |
| $K_8$ | 17.87 | 18.36 | 7.82 | 6.96 | 22.51 | 14.52 | **2.69** | 4.14 |
| $K_9$ | 3.86 | 5.82 | 1.86 | 1.35 | 5.12 | 2.26 | 1.36 | 1.76 |
| $K_{11}$ | 6.70 | 5.36 | **0.76** | 2.60 | 4.32 | 2.81 | 1.73 | 3.14 |
| $K_{14}$ | 0.46 | 0.27 | 0.05 | 0.07 | 0.65 | 0.13 | 0.09 | 0.04 |
| $K_{15}$ | 1.73 | 0.59 | **0.18** | 0.21 | 1.03 | 0.55 | 0.24 | 0.40 |
| $K_{16}$ | 4.48 | 2.60 | 1.99 | 1.75 | **0.32** | 3.97 | 2.15 | 1.76 |
| $K_{17}$ | 6.73 | 4.54 | 2.65 | 2.00 | **0.67** | 0.75 | 2.66 | 1.95 |
| $K_{23}$ | 3.26 | 3.15 | 2.43 | 2.20 | 2.97 | 1.55 | 1.84 | 2.78 |

**Table S11.** Tanh-cPIKANs: Relative parameter estimation errors (%) under fixed (F) and randomly resampled (R) collocation point configurations. F-*N* denotes training with *N* fixed collocation points, while R-*N* denotes resampling *N* points at each epoch. Errors are computed relative to nominal parameter values after convergence. Entries in **bold** indicate the most accurate configuration (lowest relative error) for each parameter across sampling strategies and Network architecture.

| Params | F-100 | F-200 | F-1000 | F-1500 | R-100 | R-200 | R-1000 | R-1500 |
| --- | --- | --- | --- | --- | --- | --- | --- | --- |
| $K_8$ | 21.05 | 15.66 | 7.16 | 6.12 | 14.00 | 14.38 | 6.45 | 8.16 |
| $K_9$ | 4.39 | 2.09 | 2.42 | **1.33** | 4.14 | 2.96 | 4.54 | 1.64 |
| $K_{11}$ | 26.01 | 15.11 | 12.50 | 7.18 | 16.17 | 11.50 | 6.39 | 4.54 |
| $K_{14}$ | 12.02 | 3.27 | 0.63 | 0.09 | 2.60 | 1.73 | 0.76 | **0.01** |
| $K_{15}$ | 20.49 | 17.28 | 5.17 | 1.24 | 5.96 | 6.37 | 3.84 | 3.27 |
| $K_{16}$ | 2.45 | 2.19 | 1.44 | 1.47 | 2.67 | 0.61 | 0.51 | 1.37 |
| $K_{17}$ | 3.36 | 2.93 | 0.73 | 1.50 | 5.16 | 0.57 | 1.65 | 1.41 |
| $K_{23}$ | 1.84 | 1.33 | 1.92 | 1.56 | 2.20 | 2.57 | **1.17** | 1.85 |

